# REpair of heterozygous Mutations independent of Exogenous Donor template with high efficiency (REMEDY) using allele specific CRISPR targeting and HDR enhancers

**DOI:** 10.1101/2025.05.29.641201

**Authors:** Yasemin Sezgin, Genesis Snyder, Noushin Saljoughian, Colin Maguire, Ebru Erzurumluoglu Gokalp, Devi Jaganathan, Eleonora S D’Ambrosio, Burcak Ozes, Gregory Wheeler, Ben Kelly, Mark Hester, Zarife Sahenk, Sriram Vaidyanathan, Afrooz Rashnonejad, Jerry Mendell, Nizar Y. Saad, Roshini S. Abraham, Juhi Bagaitkar, Susan D. Reynolds, Allison Bradbury, Meisam Naeimi Kararoudi

## Abstract

We report the development of REMEDY (REpair of heterozygous Mutations independent of Exogenous Donor template with high efficiencY), a genome editing strategy that allows efficient repair of heterozygous mutations in human and mouse cells without necessitating an exogenous donor DNA template. Here, we used in-PAM or near-PAM CRISPR strategies to induce a double-strand break (DSB) in mutant alleles. Following the DSB, the wild-type homologous chromosome itself serves as an endogenous DNA donor template and initiates the correction of the mutant allele. Concurrently treating the cells with HDR enhancers, such as AZD7648, further improved the efficiency of the correction. We demonstrated the utility of REMEDY in the context of six different diseases with heterozygous mutations such as IBMPFD, cystic fibrosis, progeria, ITPR3-associated combined immunodeficiency, ACTA1, and TBCD in human patient derived primary cells and complementary mouse model cell lines.

Graphical Abstract

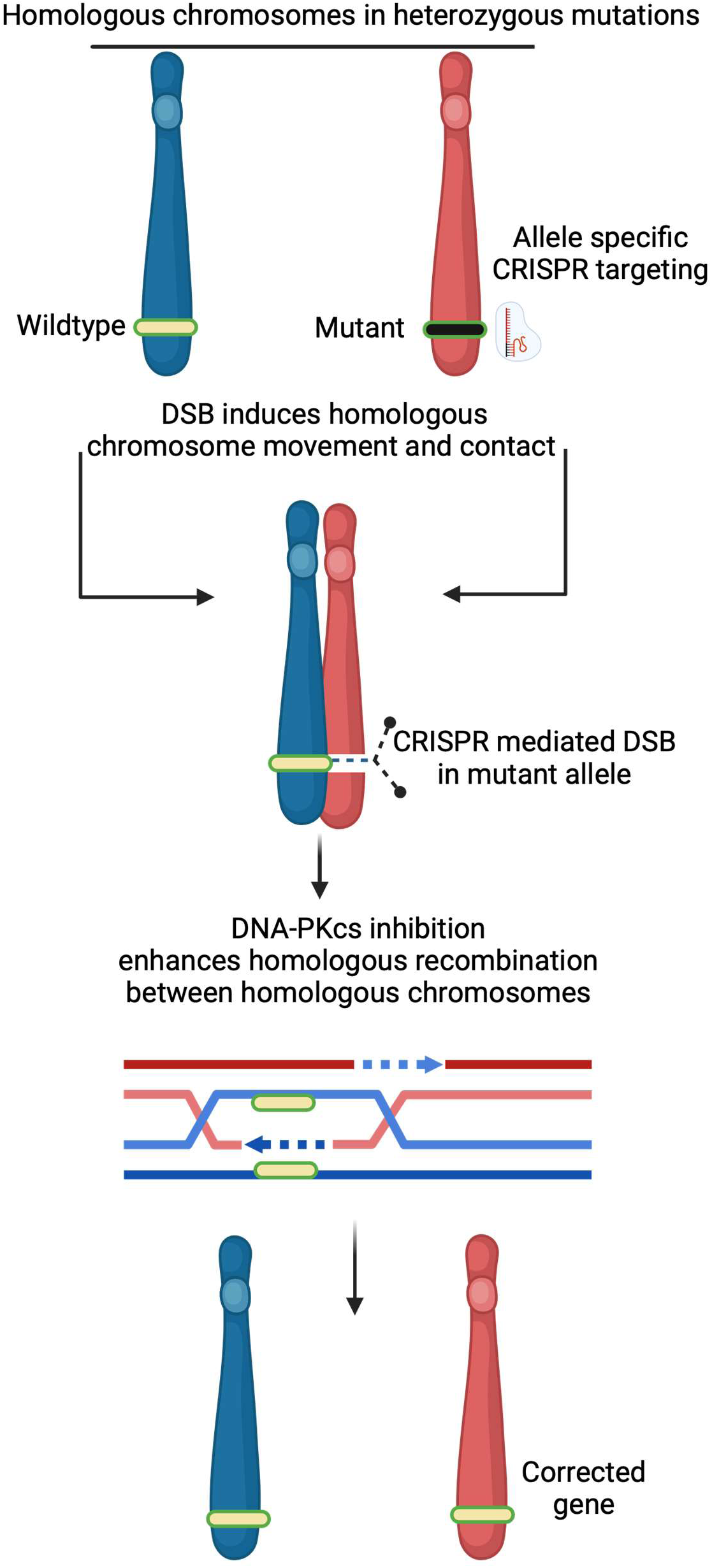

## Introduction

Heterozygous mutations affect either a single allele in dominant or sex-linked inheritance patterns, or both alleles in the case of biallelic autosomal recessive inheritance. These mutations can be corrected by clustered regularly interspaced short palindromic repeats (CRISPR) mediated homology-directed repair (HDR) and exogenous DNA templates. However, introducing exogenous HDR templates into the cells via electroporation, infection, lipofection, or by viral delivery approaches can lead to cell death, loss of regenerative potential, and adds additional manufacturing and financial burdens^1^. Therefore, there is an essential need to develop a novel approach to efficiently edit heterozygous mutations independent of an exogenous DNA donor template. Here we report REMEDY: REpair of heterozygous Mutations independent of Exogenous Donor template with high efficiencY.

In G₀-phase in human cells, double-strand breaks (DSB) result in movement of the homologous chromosomes to the DSB site and formation of a transient contact^2^. Other studies report that CRISPR-induced DSB triggers recombination between homologous chromosome arms in fly lines^3^. Finally, homologous chromosome exchange has been reported in mouse cells with biallelic mutations^4^. Therefore, we posited that we could trigger this homologous recombination mechanism between wild-type and mutant homologous chromosomes in human and mouse cells that harbor heterozygous mutations. To initiate the HDR between the homologous chromosomes, a DSB is introduced only in the mutant allele. Mutant allele-specific targeting in heterozygous mutations can be achieved by designing gRNAs that contain mutated nucleotide(s) located near-PAM (within the 3’ seed sequence of the gRNA) or in-PAM sequence^5,6^. Therefore, we hypothesized that an in-PAM or near-PAM gRNA design strategy would induce a targeted DSB only in the mutant allele and initiate the recruitment of the homologous healthy chromosome to serve itself as endogenous DNA donor template for correction of heterozygous mutations. This strategy does not require an exogenous DNA donor template for mutation correction. We called our novel approach REMEDY.

## Results

### REMEDY corrects VCP gene mutations in mouse and patient derived cells of IBMPFD

To test REMEDY, we first focused on inclusion body myopathy with early-onset Paget disease and frontotemporal dementia (IBMPFD). The disease is caused by a missense mutation in Valosin Containing Protein (*VCP*) gene that has autosomal dominant inheritance pattern. Here, we used a mouse model of inclusion body myopathy with a heterozygous mutation in the *VCP* gene (c.464_465GG>AT; p.Arg155His), one of the most frequent pathogenic variants in the *VCP* gene. We designed four mutant allele specific gRNAs (Supplementary table 1) to target the R155H mutation. While we observed no targeting in fibroblasts derived from healthy mice for gRNA2-4, gRNA1 targeted the wild-type allele (Supplementary figure 1) and thus was not used for the downstream experiments. We then tested the mutant allele specific gRNAs 2-4 in *Vcp^R155H/+^* mouse myoblasts. All three of the tested gRNAs resulted in complete (100%) targeting of mutant allele determined by Sanger sequencing (Supplementary figure 2). As we hypothesized, we observed a modest correction of the mutant allele without an exogenous DNA template. The frequency of wild-type allele in the cells targeted by gRNA2 increased from normalized 50% to 58% as measured by next generation sequencing (NGS) and analyzed by CRISPResso2^7,8^. Therefore, 8% of the cells were corrected to homozygous for the wild-type allele without an exogenous donor DNA template. Sanger sequencing electropherograms showed the highest frequency of wild-type nucleotides at the position achieved by the gRNA2 among all the other gRNAs tested (Figure 1A and Supplementary figure 3). This data confirms that ICE (Sanger) and CRISPResso2 (NGS) analysis provide comparable results as reported before^7^.

**Figure 1.**
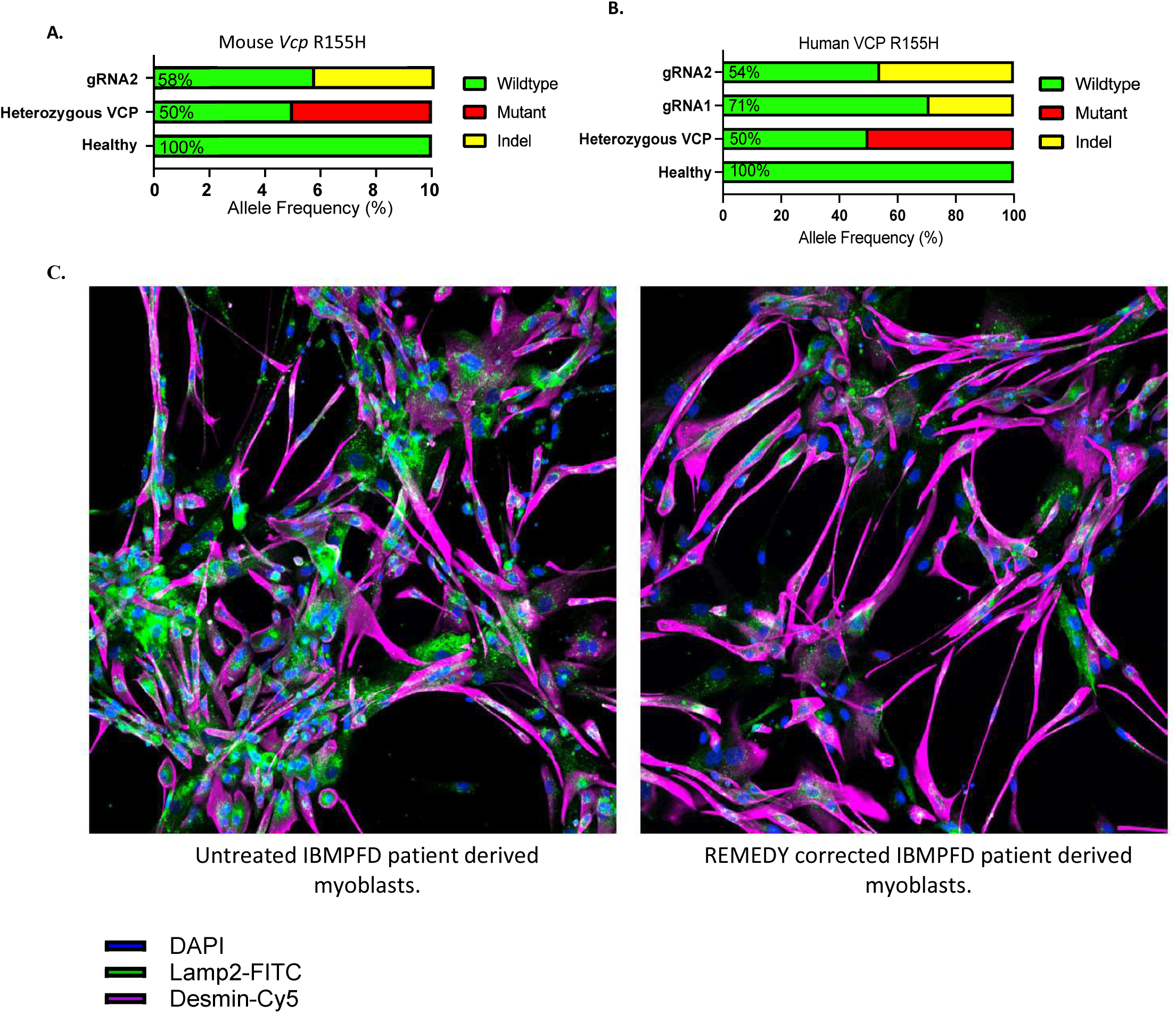
REMEDY corrected heterozygous mutations in *VCP* mouse myoblasts. (A), and in human IBFMD patient derived skin fibroblast measured by NGS analyzed by CRISPResso2 (B).LAMP2 immunostaining images of untreated and REMEDY corrected heterozygous mutations in VCP patient iPSC derived myoblasts. (C)

The VCP mouse model has different base pair substitution than the human mutation (c.464_465GG>AT in mouse equal to c.464G>A in human) coding the same amino acid. To ensure that the REMEDY is not mutation and cell type specific, we then designed two gRNAs and targeted the human *VCP* heterozygous mutation c.464G>A (p.Arg155His) mutant allele and tested these guides in IBMPFD patient derived fibroblasts. Sanger sequencing data showed an increase in the frequency of wild-type allele from 50% to 71% with gRNA1 and to 54% with gRNA2 without using an exogenous donor DNA template (Figure 1B and Supplementary figure 4A). To ensure that the REMEDY is not pathogenic allele specific, we targeted the wildtype allele of *VCP* c.464G with a gRNA that was designed specific to wild type allele and tested the guide in patient derived myoblasts and generated the homozygous mutation of *VCP* c.464G>A (p.Arg155His) with high efficiency (59% mutant allele) as analyzed by CRISPResso2 (Supplementary figure 4B and Supplementary table 2).

### Homology directed repair enhancers increase the efficiency of REMEDY

The homology repair pathway takes place during the S and G2 phases when a DNA template is available such as a sister chromatid or an exogenous DNA template. It has been shown that HDR enhancers, such as IDT HDR enhancer and AZD7648, arrest or pause the cell cycle at G1 and S at high frequency^9^. Since the REMEDY correction is regulated by HDR between homologous chromosomes, we then hypothesized that the addition of HDR enhancers such as IDT Alt-R HDR Enhancer V2 and AZD7648 as a DNA-PK inhibitor may further improve the efficiency of correction achieved by REMEDY^10^. Here, we report that AZD7648-mediated cell cycle synchronization favors homology repair between homologous chromosomes and increases the correction of the mutant allele. However, the exact mechanism remains to be verified.

IBMPFD effects muscles. Therefore, we acquired patient iPSC derived myoblasts and tested REMEDY on these cells and observed an increase in the wild-type allele frequency from 50 to 69% using gRNA2 with AZD7648 (Supplementary Figure 5). To evaluate whether this genetic correction translated into phenotypic improvement, we performed immunostaining with antibodies against lysosome-associated membrane protein (LAMP2), a well-established marker of autophagy and observed a partial reduction in LAMP2 expression (Figure 1C). It is worth mentioning that further functional assays are required.

### REMEDY corrects heterozygous mutations in tubulin folding cofactor D (*TBCD*) gene in patient derived fibroblasts

We further tested REMEDY in the context of tubulin folding cofactor D (*TBCD*) in human patient-derived fibroblasts. Biallelic autosomal recessive mutations in this gene result in a rare early-onset encephalopathy with neurodevelopmental and neurodegenerative features. Skin biopsy samples were collected from two different patients, patient 1 mutation 2 (P1M2) (TBCD; c.2305-2307 del GAG p.E769del) and patient 8 mutation 2 (P8M2) (*TBCD*; c.2472-2A>G-intronic splice acceptor mutation). Specifically, we designed allele specific gRNAs for the mutant allele (Supplementary table 3 and 4) and patient derived fibroblasts were electroporated with Cas9/RNP targeting the mutant allele. In patient P1M2, we observed an increase in percentage of wild-type allele in the cells targeted by the mutant allele specific gRNA3 from 50% to 64% (Figure 2A), and in P8M2, REMEDY achieved 8% correction (Supplementary figure 6A and B) as measured by ICE and CRISPresso2. In P1M2, treating the cells with IDT Alt-R Enhancer V2 or AZD7648 post Cas9/RNP electroporation resulted in a further increase in the frequency of the wild-type allele to 83% and to 82% in the cells, respectively, as measured by ICE. This means 30% of the cells have become homozygous for wild-type TBCD. We then further performed NGS and analyzed the data with CRISPResso2 and identified similar percentage of correction (22% wild-type homozygous *TBCD*)(Figure 2B and C). The NGS and Sanger sequencing on P8M2 also showed enhanced correction after treating the cells with AZD7648 and IDT HDR enhancer (from 50% to 70%) along with REMEDY (Supplementary figure 6A and B).

**Figure 2.**
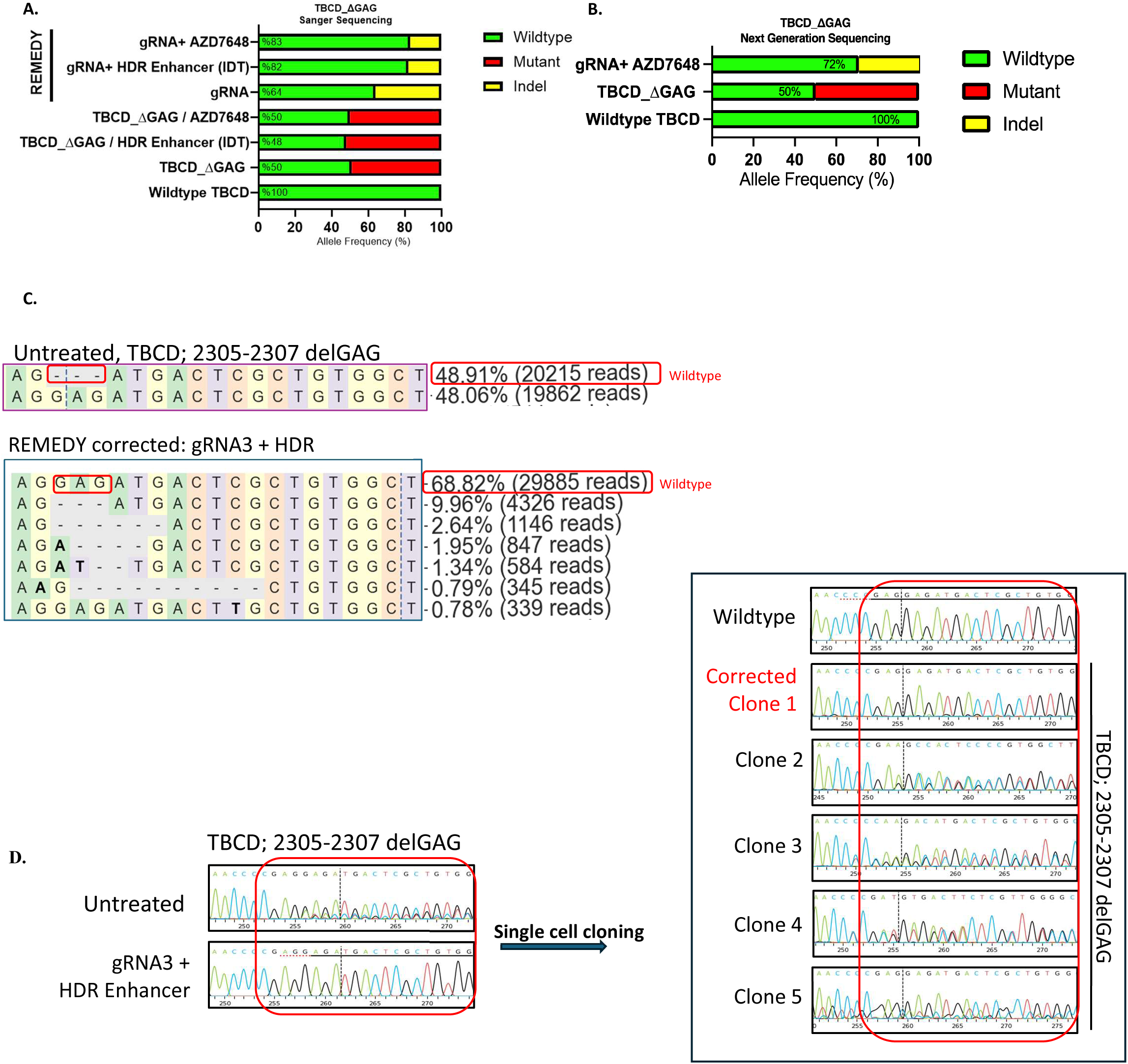
REMEDY corrected mutation in *TBCD* gene and the addition of HDR enhancer enhances the correction measured by Sanger sequencing P1M2 using gRNA3. (A). NGS demonstrated similar correction rate detected by Sanger sequencing (B and C). Single cell cloned REMEDY edited TBCD fibroblasts confirmed the sequencing results and showed the edits are stable after growing cells for 10 days (D).

### Single cell clones of the REMEDY corrected cells confirmed the correction of heterozygous mutation in TBCD

To ensure the stability of the edits and to confirm that our findings are not related to sequencing artifacts, we performed single cell cloning of the patient derived fibroblasts that were targeted with allele specific gRNA3 and treated with HDR enhancer. Based on Sanger sequencing and NGS data we expected to find one out of five clones (20%) to be fully corrected to wild-type homozygous *TBCD*. This was confirmed by the Sanger sequencing of five single cell clones that were grown for more than 10 days (Figure D).

### REMEDY corrected heterozygous mutation in ITPR3 gene

We tested REMEDY on a heterozygous mutation in the inositol 1,4,5 triphosphate receptor 3 (*ITPR3*) which regulates intracellular calcium stores, and inherited pathogenic variants can cause Charcot-Marie-Tooth disease, demyelinating, type 1J and a severe combined immunodeficiency^11^. We obtained patient-derived fibroblasts and designed a gRNA (Supplementary table 5) to target only the *ITPR3* allele harboring the mutation (c.7570C>T, p. Arg2524Cys). REMEDY, with the addition of HDR enhancers, again was able to correct the heterozygous mutation in these cells with no need for exogenous DNA donor template, normalized 50% to 62% with AZD7648 and 50% to 61% with IDT HDR enhancer V2 measured by NGS and analyzed by CRISPResso2 (Figure 3A and B) and Sanger following analysis by ICE (Supplementary figure 8).

**Figure 3.**
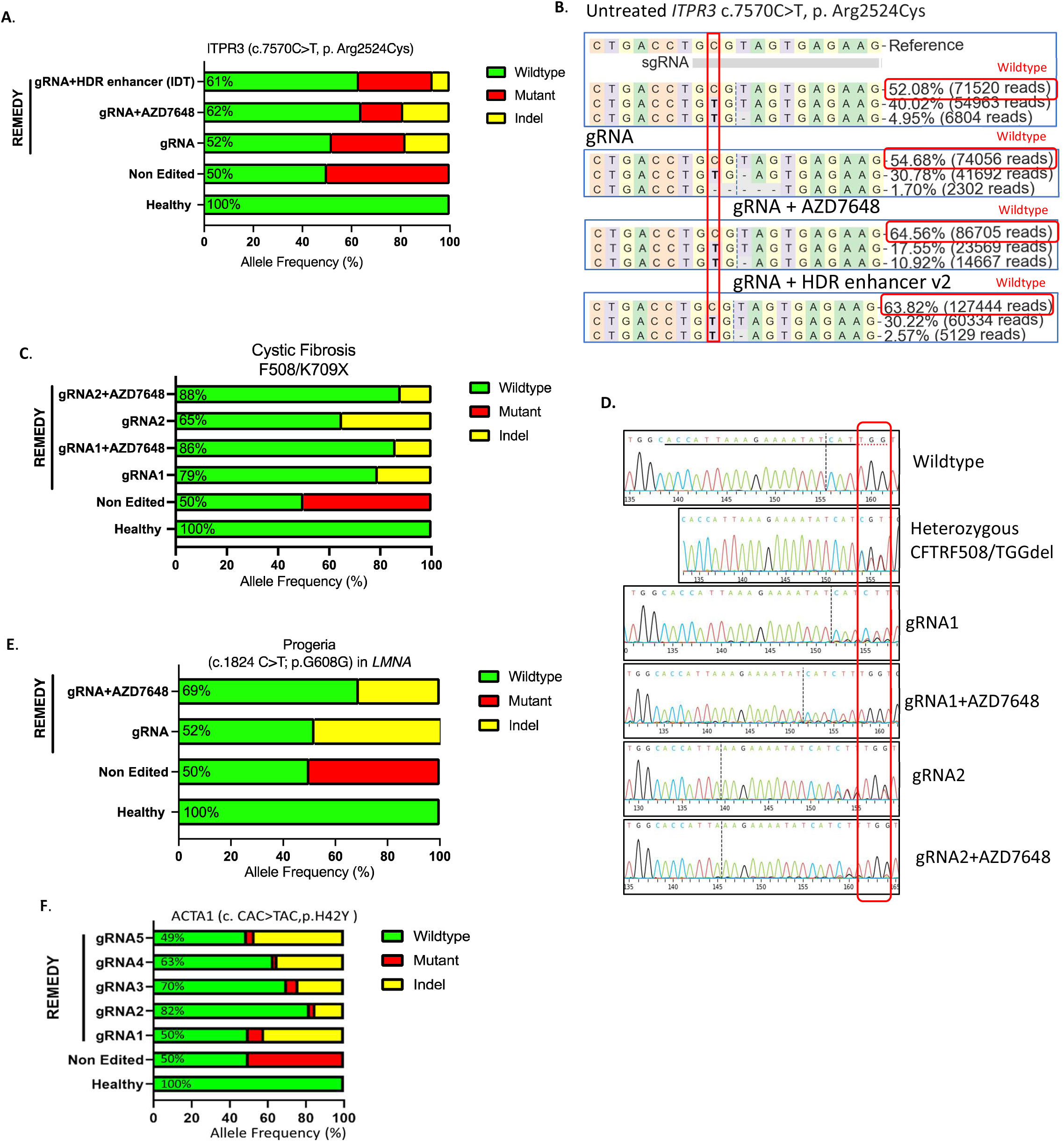
REMEDY corrects several heterozygous mutations in different cell types and diseases. REMEDY corrected heterozygous mutations in *ITPR3* heterozygous mutation in human skin fibroblast measured by NGS (A and B); and in human airway epithelial cells with *CFTR* heterozygous mutation derived from a patient with CF measured by Sanger sequencing (C and D). We observed similar results in fibroblasts derived from Progeria patients measured by Sanger sequencing (E). REMEDY corrected mutation in ACTA1 gene measured by Sanger sequencing (F)

Since AZD7648 and IDT HDR enhancer V2 resulted in comparable efficacy and AZD7648 is used in clinical settings, we performed the rest of the experiments using only the AZD7648 DNA-PK inhibitor. It worth mentioning despite achieving corrections, the *ITPR3* pathogenic variant appears to confer a survival disadvantage to fibroblasts, compared to healthy control fibroblasts or those from patients with other inborn errors of immunity (IEIs; e.g. IPEX). Further functional studies are required.

### Remedy corrected the most common heterozygous mutation in CFTR gene in CF patient derived bronchial epithelial cells

To test whether REMEDY could be used for *ex vivo* cell therapy applications, where clinical implications could be transformative, we used human airway epithelial cells collected from a patient with cystic fibrosis (CF). We designed allele-specific gRNAs targeting a deletion of phenylalanine 508 (F508del) in *CFTR* gene (Supplementary table 6)^12^. A single heterozygous mutation impairs CFTR folding resulting in chloride channel dysfunction^12^. We electroporated human bronchial epithelial cells (HBEC) harboring one copy of the F508del variant with the Cas9/RNP, treated the cells with AZD7468, and observed a significant increase in the frequency of wildtype allele in the cells corrected with gRNA1 from 50% to 86% and to 87% with gRNA2. Confirming the potential of REMEDY for genome editing free of an exogenous donor DNA template in a cell type that has therapeutic implications for ex vivo cell therapy (Figure 3C and D).

### REMEDY results in correction of human derived cells from Hutchinson–Gilford progeria syndrome (HGPS or progeria)

Hutchinson–Gilford progeria syndrome (HGPS or progeria) is caused by a dominant-negative mutation in the *LMNA* gene (c.1824 C>T; p.G608G) which encodes nuclear Lamin A and Lamin C^13^. In previous studies, base editing corrected this mutation in patient-derived fibroblasts with an efficiency of 87–91% of the pathogenic allele^6^. Here, we used REMEDY to correct the same mutation in Progeria patient-derived fibroblasts and achieved highly efficient targeting of the mutated allele (100%) and achieved correction of the mutated allele (50% to 69% wild-type allele) in the cells treated with AZD7648 post allele specific CRISPR targeting without an exogenous DNA template (Figure 3E and Supplementary table 7).

### REMEDY corrected the autosomal dominant mutations in the skeletal muscle alpha (α)-actin (*ACTA1*) gene in iPSC cells

We tested the REMEDY approach to correct autosomal dominant mutations in the skeletal muscle alpha (α)-actin (*ACTA1*) gene in patient derived iPSC cells. *ACTA1* is the primary actin isoform found in skeletal muscles and thin filaments in the sarcomere, the basic unit of muscle contraction ^14,15^. Mutations in this gene are linked to disrupted actin polymerization and impaired interactions with other proteins such as myosin^16–18^, ultimately leading to a severe Nemaline myopathy characterized by muscle weakness, delayed developmental milestones, respiratory difficulties, and dilated cardiomyopathy ^14,15,19–21^. For this experiment, we designed allele-specific gRNAs targeting the H40Y mutation in the ACTA1 gene (p. His42Try, c. CAC>TAC, 124C>T)(Supplementary table 8). We applied REMEDY in H40Y ACTA1 iPSC cells (generous gift from Dr. David Mack, University of Washington)^22^ and analyzed with Sanger sequencing. Sequencing results demonstrated 82% correction of the mutant allele using gRNA2 while the wild-type allele remained unaffected (Figure 3F). These results confirm the efficacy of REMEDY in editing autosomal dominant mutations in *ACTA1* without requiring an exogenous HDR template. This is important since many pathogenic mutations in the *ACTA1* gene have been associated with NEM3 disease and having a simple and cost-effective gene editing platform will be crucial toward advancing therapeutic approaches. Furthermore, since there are six actin genes in human cells with ∼90% sequence homology, using REMEDY ensures a highly specific genome-editing approach without relying on HDR templates to minimize the risk of off-target effects.

### PacBio sequencing did not show evidence of large-scale genomic alterations

Cas9 mediated large DNA resection resulting in large-scale genomic alterations such as loss of heterozygosity (LOH) has been a concern surrounding the use of this technology ^23–25^. Additionally, DNA-PK inhibition may further increase the frequency of the LOH^26^. Therefore, we performed PacBio HiFi long-read sequencing to evaluate such potential alterations after the use of REMEDY at the TBCD-P1M2 pathogenic allele target site (chr17: 82,924,980, TBCD; c.2305-2307 delGAG p.E769delP1M2) in untreated and REMEDY corrected patient derived iPSC cells. PacBio HiFi sequencing was performed and the data were aligned to the human reference genome (GRCh38) using pbmm2 (v1.13.1) (Supplementary Table 10) ^27,28^ ^29^. Overall, PacBio indicated no meaningful LOH effects near the target site.

In detail, manual inspection of reads at the target site confirmed the expected variant in the untreated sample in the heterozygous condition (Figure 4A). In the REMEDY corrected sample, we showed the high efficiency of targeting the mutant allele as shown by Sanger sequencing and NGS (18 delGAG reads untreated vs. 2 delGAG reads in REMEDY corrected).Therefore, two delGAG reads remained, as well as 5 reads containing short non-frame-shifting deletions of uncertain pathogenicity. The remaining altered reads contained either short frame-shifting deletions (8) or large deletions (2). A corrective effect is expected in 58-88% of the altered alleles observed.(Figure 4B and C, Supplementary Table 9 A and B).

**Figure 4.**
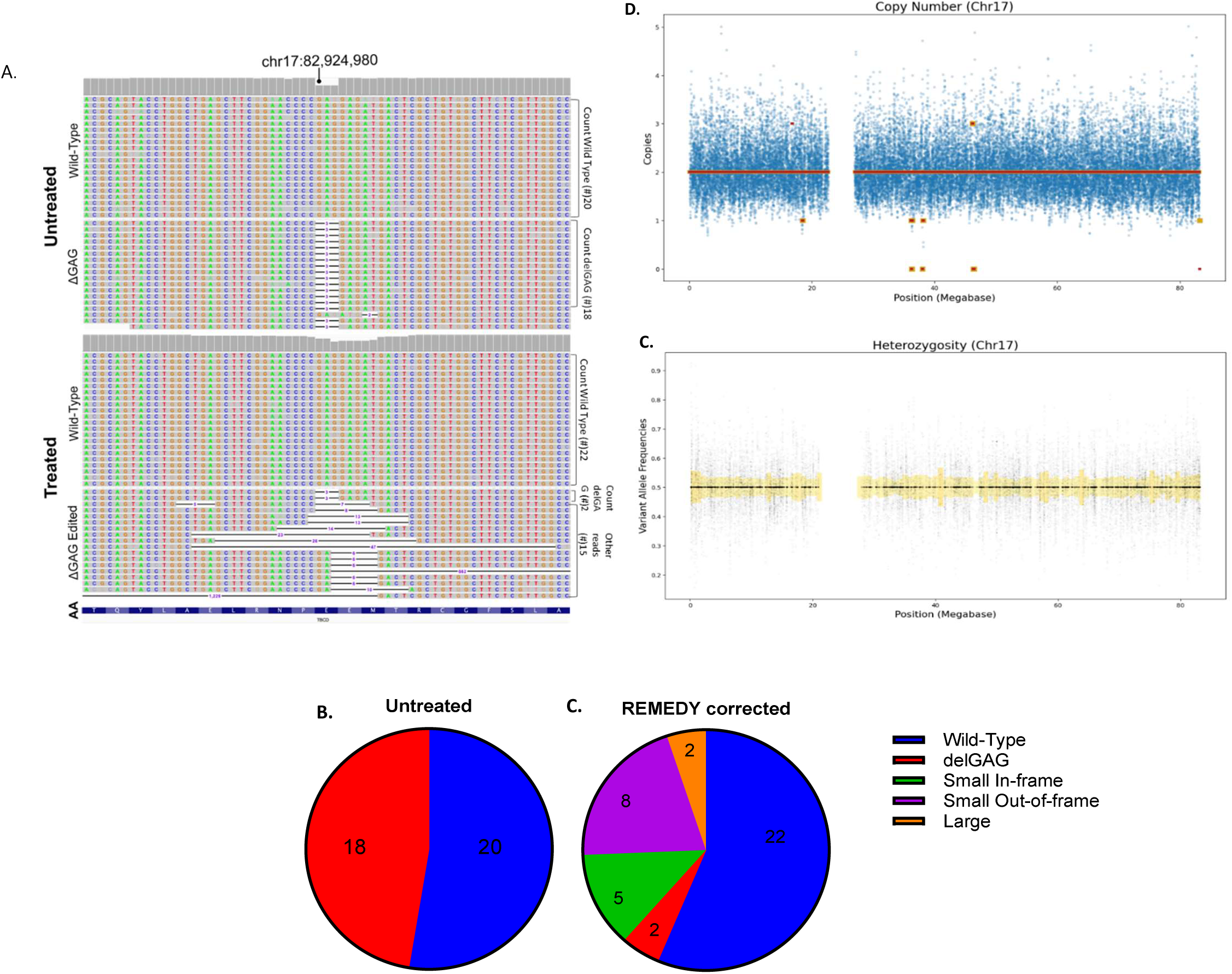
Visualization of Untreated and Post-REMEDY Corrected Samples in IGV. While the untreated sample shows only two alleles (wild-type and delGAG), the REMEDY corrected sample exhibits replacement of most delGAG alleles, instead showing an assortment of other deletion sizes. The in-frame amino acid (AA) sequence for the gene TBCD at the position of interest is shown below (A). **Allele composition in Untreated and Post-REMEDY corrected samples.** Untreated **(B)** sample shows a heterozygous state of the wild-type and delGAG alleles. REMEDY corrected **(C**), most delGAG allele reads are replaced with deletions of other sizes. **Copy-Number Analysis of Chromosome 17.** Blue background points show copy number calculated at a given location from post-REMEDY correction coverage depth, normalized against coverage of the untreated sample. Yellow and red segments show the HiFiCNV-calculated copy state at a given position for the pre– and REMEDY corrected samples, respectively. Overlapping red and yellow segments show copy states which are consistent between both samples (D). **Loss-of-Heterozygosity Analysis of Chromosome 17.** Black points show variant allele frequency called at a given position. Yellow bars show allelic spread (symmetric distance of calculated average from 0.5) for regions each representing 1% of the chromosome(E) demonstrating that there is no detectable LOH.

DeepVariant analysis detected 33 phased heterozygous sites within 10kb of the target site in the untreated sample. In the corrected sample, all sites remained heterozygous; only one (chr17: 82932762) showed any discrepancy, with 2 of 38 (5.3%) reads deviating from the expected allelic phasing. This represents only 0.2% of total reads and is unlikely to indicate any meaningful LOH effects near the target site. (Supplementary Table 10)

No large-scale copy-number changes were observed in chromosome 17. Nine small focal gains or losses were detected, of which seven matched to the untreated comparator. One additional single-copy gain was detected on 17p (chr17:16752000-16844000) and a single-copy loss at the distal end of 17q (chr17:83234000-83256000) was instead called as a complete loss (Figure 4D); however, manual inspection of these regions in Integrative Genomics Viewer (IGV) showed no discernible differences between the corrected and untreated samples. We confirmed the degree of coverage variation observed across chromosome 17 was comparable to chromosome 12 (Supplementary Figure 9).

After filtering for appropriate coverage and appropriate allele frequency in the untreated comparator sample, 57481 variants were used to determine post-treatment loss-of-heterozygosity. Chromosome 17 heterozygosity bins deviated from the expected variant allele frequency (VAF) by 2.2% to 6.7%, comparable to the deviations observed in chromosome 12 (1.2% to 5.6%) (Figure 4E and Supplementary Figure 10).

## Conclusion

REMEDY has the potential to be used in basic science and pre-clinical studies for which a heterozygous mutation needs to be corrected. It can also be potentially used to generate pathogenic homozygous mutations for *in vitro* studies if only wild-type allele is targeted, and the mutated homologous chromosome is used as the endogenous HDR template. Additionally, REMEDY has potential to treat patients with a wide variety of heterozygous mutations, especially if the correction can be performed *ex vivo* for cell therapy applications.

REMEDY has several advantages over existing gene correction approaches. In the pre-clinical setting, high efficiency of correcting heterozygous mutations can be achieved by introducing donor DNA template to the cells; however, this can be cytotoxic given the initiation of DNA sensing mechanism. In the clinical setting, providing an exogenous DNA template is expensive and not widely accessible especially in low-middle income countries.

Ongoing studies are assessing the range of correction that can be achieved by REMEDY, particularly in the context of different mutation types (single nucleotide polymorphisms vs. deletions of varying sizes. The allele specific gRNA approach used in REMEDY is another advantage of the method as it potentially minimizes the number of off targets since it is designed to only target the mutant allele. However, the in-PAM or near-PAM gRNA design can also be a limitation if there are no PAM sequences in close vicinity of the mutation site. This problem can be potentially solved by changing the type of endonuclease that expands the range PAM sequences to initiate a DSB in the mutant allele. Tools such as AlleleAnalyzer (https://github.com/keoughkath/AlleleAnalyzer) can be used for allele-specific sgRNA design and for identifying potential endonucleases^30^. Lastly, the addition of HDR enhancers improves efficiency of REMEDY. The direct comparison of two relevant enhancers validates this confirmation and enhances translation of our novel REMEDY approach.

## Short summary

Herein, we demonstrate a new mechanism that overcomes many current barriers of gene editing by efficiently repairing heterozygous mutations without the necessity of exogenous donor DNA template. We confirm these findings in 6 diseases, 2 species, and 3 cell types in the context of IBMPFD, TBCD, CF, Progeria, ITPR3, and ACTA1 which result from distinct molecular mechanisms, demonstrating the potential breadth of application of this technology.

## Methods

### Progeria Patient Derived Fibroblasts

This study was performed in compliance with the standards set by the National Institute of Health (NIH) and was reviewed and approved by the Institutional Review Board (IRB) at Nationwide Children’s Hospital (IRB number 14-00719). Written, informed consent was obtained from the participants prior to inclusion in the study. Samples from the participants were identified by numbers, not names. Patient derived human skin fibroblasts were collected under IRB and maintained in Dulbecco’s modified Eagle’s medium (DMEM, Gibco™, Catalog # 11960044) supplemented with GlutaMAX (Gibco™, Catalog #35050061) and 15% heat inactivated Fetal Bovine Serum (GenClone Catalog # 25-514H) FBS at 37°C and 5% CO_2_. Human primary Progeria dermal fibroblast cell lines were obtained from The Progeria Research Foundation (PRF) Cell and Tissue Bank. The HGPS cell lines were HGADFN367. Progeria fibroblasts were grown in DMEM media supplemented with GlutaMAX, 20% FBS, 1% Non-Essential Amino Acids (NEAA) (Thermo Fisher, Catalog # 11140050) and 1% Pen/Strep (Gibco™, Catalog # 15070-063).

### Mouse myoblasts and fibroblasts

All animal experiments were performed in compliance with the standards set by the National Institute of Health (NIH) and were reviewed and approved by the Research Institute at Nationwide Children’s Hospital Animal Care and Use Committee (IACUC approval number: AR18-00123). All experiments were conducted in compliance with the ARRIVE guidelines. All mice were sacrificed in accordance with ethical standards; overdose of xylazine/ketamine anesthesia was used to euthanize the mice.

Skeletal muscles were collected from one *Vcp^R155H/+^* mouse after euthanasia (The Jackson Laboratory, Strain #:021968). Protocol was adapted from Shahini et al., 2018^31^. Shortly, muscles were minced and seeded on a Matrigel (Corning, Catalog #354234) coated cell culture dish in proliferation medium for release of myoblasts (high glucose DMEM, 20% FBS (Thermo Fisher, Catalog #16000044), 10% horse serum (ThermoFisher, Catalog #26050070), 0.5% chicken embryo extract (Fisher Scientific, Catalog #NC9997754), 2.5 ng/ml bFGF (Peprotech, Catalog #450-33), 10 μg/ml gentamycin (Gibco, Catalog #15-710-064), 1% Antibiotic-Antimycotic (ThermoFisher, Catalog #15240062), and 2.5 μg/ml plasmocin prophylactic (Invivogen, ant-mpp). Released cells were pre-plated to purify myoblasts. Similarly, skin sample of one *Vcp^R155H/+^* mouse was collected after mouse is euthanized and fibroblasts were released starting in one week. Cells were maintained as mentioned in the Patient-Derived Fibroblasts section. All animal experiments were performed according to the ethical guidelines approved by The Research Institute at Nationwide Children’s Hospital Animal Care and Use Committee (IACUC approval number: AR18-00123).

### Airway Epithelial cells

Human bronchial epithelial cells (HBEC) were obtained from CF donor lungs through the Epithelial Cell Core at Nationwide Children’s Hospital. HBECs were cultured using Pneumacult-Ex Plus media with ROCK inhibitor (Y-27632) at 10 µM. Five days after seeding, the cells were dissociated by treatment with TrypLE and resuspended in OPTI-MEM (Gibco™, Catalog #31985070) at a concentration of 5 million cells/ml. 6 µg of Cas9 and 3.2 µg of sgRNA were mixed and incubated for 10 minutes. 20 µL of cells in OPTI-MEM were added to the Cas9/sgRNA mixture and electroporated using a Lonza 4D nucleocuvette strips. The program CA-137 with buffer setting P3 (Lonza, Catalog# V4XP-3032) was used. Five days after editing, genomic DNA was isolated from HBECs. Exon 11 locus of CFTR was amplified using PCR with an annealing temperature of 58°C and extension time of 35 sec using Q5 polymerase.

### Patient derived iPSCs

To generate patient derived iPSCs, peripheral blood mononuclear cells (PBMCs) were isolated under approved IRB. The PBMCs were then converted to iPSCs using Sendai virus reprogramming kit (Thermo Fisher, A16518) and maintained in supplemented STEMFLEX™ Medium (Thermo Fisher, A3349401) on plates coated with Vitronectin (VTN) Recombinant Human Protein, Truncated, 10mL (Thermo Fisher, A31804) at 37°C and 5% CO_2_.

### Single Cell Cloning

Cells were detached and diluted to 1×10^4^ cells/ml in culture media. A suspension of 5 cells/mL was prepared from the 1×10^4^ cells/mL solution by adding 25 µL to 50 mL of culture media. 200 µL of the new dilution was added to each well of a 96 well plate. The plates were monitored for 7-10 days to assess for single cell colonies. Upon reaching an appropriate cell number, PCR was performed on the isolated DNA, purified, and sent for Sanger sequencing to validate a homogeneous population.

### Preparation of Allele specific gRNAs

*VCP*, *TBCD*, *CTFR*, *ACTA1*, *LMNA* and *ITPR3* gRNAs were designed by using online tool Benchling (https://benchling.com) and synthesized by Synthego as Synthetic gRNAs resuspended in TE buffer to achieve 100 µM working concentration.

### Performing allele specific CRISPR in mouse myoblasts and patient derived fibroblasts, iPSCs and myoblasts

Fibroblast cells were detached at least 70% confluency using TrypLE Express Enzyme (Fisher Scientific, Catalog #12-604-021). The cells were collected and counted using trypan blue. A total of 1.0×10^5^ cells were used for electroporation. The cells were electroporated using the Cas9/RNP as 2 μl of Alt-R™ S.p. Cas9 Nuclease V3, 500 µg (IDT, Catalog # 1081059) at 62 μM, 2 μl gRNA, and 1 μl of DPBS (Corning, Catalog # 21-031-CV). The control cells received 5 µL of DPBS. the prepared Cas9/RNP was incubated at RT for 20 minutes. The cells were resuspended in 20 μl SE electroporation buffer (Lonza, Catalog #V4SC-1096). Resuspended cells were combined with the cas9/RNP and loaded into the electroporation cuvette provided in the Lonza kit. The cuvette was placed into the Amaxa 4D-Nucleofector X Unit (Lonza, Catalog # AAF-1003X). The cells were electroporated with the parameters of SE buffer and a pulse code of CD-137. Post electroporation, the cells were rested for 1 minute at RT, then using prewarmed culture media the cells were transferred to 12 well plates containing the prepared media. Cells recovered in the incubator at 37°C, 5% CO_2_. hiPSCs derived myoblasts were obtained from VCP donor patients through the Cure VCP Disease Foundation. Cells were seeded onto 10 cm culture dishes coated with Collagen (Col I, Corning, Cat. 354236) and grown in Myoblast expansion medium (iXCells, Catalog # MD-0102A1) with 1% Pen/Strep (Gibco™, Catalog # 15070-063). Medium was changed every other day and cells were passaged once they reached 80-90% confluency using TrypLE (Gibco™ Catalog # 12604021). Human and mouse myoblasts were treated in a similar fashion with the only variation being 0.25×10^5^ cells were used per electroporation. Electroporation on iPSCs was performed using P4 electroporation buffer (Lonza, V4XP-4032) and a pulse code CA-137. Following electroporation, cells were cultured in supplemented STEMFLEX™ Medium (Thermo Fisher, A3349401) with 1X CultureSureTM CEPT Cocktail(1,000×) (FujiFilm, 033-26071). Medium was changed 24-hours post electroporation. The medium was replaced with supplemented STEMFLEX™ Medium (Thermo Fisher, A3349401), without antibiotics, containing 1X Y27632 Rock Inhibitor (LC Labs, Y-5301). To enhance HDR we treated the post electroporation cells with 1 mM solution of AZD7648 (R&D Systems, Catalog # 7825/10) was prepared in DMSO. 1 μL AZD7648 was added per 1 mL of media. The IDT HDR enhancer V2 was used at concentration of 10 μM. The media containing AZD7648 and IDT HDR enhancer V2 were removed and replaced with fresh media 24 hours after the addition of the cells.

### DNA isolation and PCR

Genomic DNA was isolated from REMEDY corrected and untreated cells using DNeasy Blood & Tissue Kit (Qiagen, Catalog #69504). Isolated DNA was subjected to PCR amplification using the Platinum™ SuperFi II PCR Master Mix (Thermo Fisher, Catalog # 12368010). Following PCR, each sample was purified following the Qiagen QIAquick PCR Purification Kit (Qiagen, Catalog # 28104). Purified samples were run on Invitrogen™ E-Gel™ Agarose Gels with SYBR™ Safe DNA Gel Stain, 2% (Fisher Scientific, Catalog # A45205) using the E-Gel™ Power Snap Electrophoresis System Starter Kit, SYBR Safe 2% (Thermo Fisher, Catalog # G8322ST) to validate the PCR product.

All PCR primers-utilized for NGS and Sanger sequencing are described in Supplementary tables 11 and 12.

### Sanger Sequencing

Purified PCR samples along with the corresponding primers were submitted to Azenta/Genewiz (www.genewiz.com) for Sanger sequencing. The received.ab1 files were analyzed using ICE (ice.synthego.com). Briefly, healthy samples were used as the control sample for each experiment. The gRNA used during a KO needs to be changed to the WT sequence without the mutation present in the patient sample.

### NGS Deep Sequencing

Purified PCR samples were sent to the Center for Computational & Integrative Biology at Massachusetts General Hospital (dnacore.mgh.harvard.edu). Samples were run on the Illumina MiSeq with 2×100bp reads. Data was analyzed through the CRISPResso2 (crispresso.pinellolab.org/) data analysis tool. Briefly, the tool aligns each read to all allelic variants present in the controls and matches the read to the allele that is more similar.

### PacBio Sequencing

#### Allele Assessment

Reads at the target site were visualized using the Integrative Genomics Viewer (IGV) and manually tallied ^32,33^. Each read was assigned to one of the following categories: wild-type, delGAG, small non-frame-shifting deletion (<100 bp, multiple of 3bp), small frame-shifting deletion (<100 bp, not a multiple of 3bp), and large deletion (>=100bp).

#### Local Loss-of-Heterozygosity

To detect the presence of local heterozygosity potentially introduced around the edit-site, phased variant-calling was performed via DeepVariant (v1.6.0) on both samples^34^. Phased heterozygous sites within 10kb of the target site were identified from the untreated sample. Findings at these sites were then analyzed in the treated sample, to detect any deviation from expected heterozygosity or disruption of allelic phasing.

#### Chromosomal CNV-LOH

To detect large-scale loss-of-heterozygosity, DeepVariant results were compared across the length of chromosome 17. Sites with a VAF between 0.40 and 0.60 were identified in the untreated sample and filtered to those with at least 10 reads of support and not within 1 megabase of a centromere region. VAFs were then compared with the corrected sample, accounting for the degree of error observed in the untreated comparator. These VAFs were then binned into equal regions each representing 1% of the chromosome’s length, averaged, and plotted. Post-treatment allele frequencies of sites found to be heterozygous in the comparator were plotted as supporting data.

HiFiCNV (v1.0.0) was used to detect the presence of CNV events^35^. The resulting copy-number estimates were plotted for both corrected and untreated samples. Coverage metrics gathered by HiFiCNV were plotted as background supporting data. Identical analyses were also performed for non-targeted chromosome 12 for comparison.

### Immunofluorescence staining

To detect the co-localization of Lamp2 and Desmin, iPSC derived myoblasts were transferred on coverslips, fixed in 4% paraformaldehyde, permeabilized with 0.1% Triton X-100 in PBS, and blocked with 1% bovine serum albumin with 10% normal goat serum and 0.1% Triton x-100. The cells were incubated with primary antibodies against LAMP2 in 1:100 dilution (Santa Cruz Biotechnology, Catalog # sc-18822) and Desmin in 1:200 dilution (Invitrogen, Catalog # MA5-16357) for 90 minutes. Afterwards, the cells were incubated with FITC (Thermo Fisher, Catalog# A11017) and Cy5 (Thermo Fisher, Catalog # A21244)-conjugated secondary antibodies in 1:500 dilution for both for 45 minutes and treated in ProLong Gold Antifade Reagent with DAPI (Invitrogen, Catalog# P36935).

## Acknowledgement

The Cure Cystic Fibrosis Columbus (C3) Epithelial Cell Core (ECC) at Nationwide Children’s Hospital and The Ohio State University provided primary human *bronchial* epithelial cultures for this work. C3 is supported by a Research Development Program Grant (MCCOY17R2) from the Cystic Fibrosis Foundation. We also acknowledge, Joshua Robinson, Mustafa Bilal Bayazit, Jingting Zhu, and Neel Lachhwani for their help in conducting some experiments. We also acknowledge CRISPR/Gene Editing and Institute of Genomic Medicine Genomic Services cores at Nationwide Children’s Hospital. We would also like to thank Dr. Dario Palmieri for his valuable comments. We also thank the Cure VCP Disease foundation that provided primary myoblasts. We would like to thank Dr. David Mack at the University of Washington for generously gifting the ACAT1 H40Y iPSC cells. The ACTA1 REMEDY experiments has been supported by NCH start-up funds to Dr. Rashnonejad.

## Data availability statement

The datasets used and/or analyzed during the current study are available from the corresponding author on reasonable request.

## Supplementary Figures and Tables

**Supplementary Table 1.**
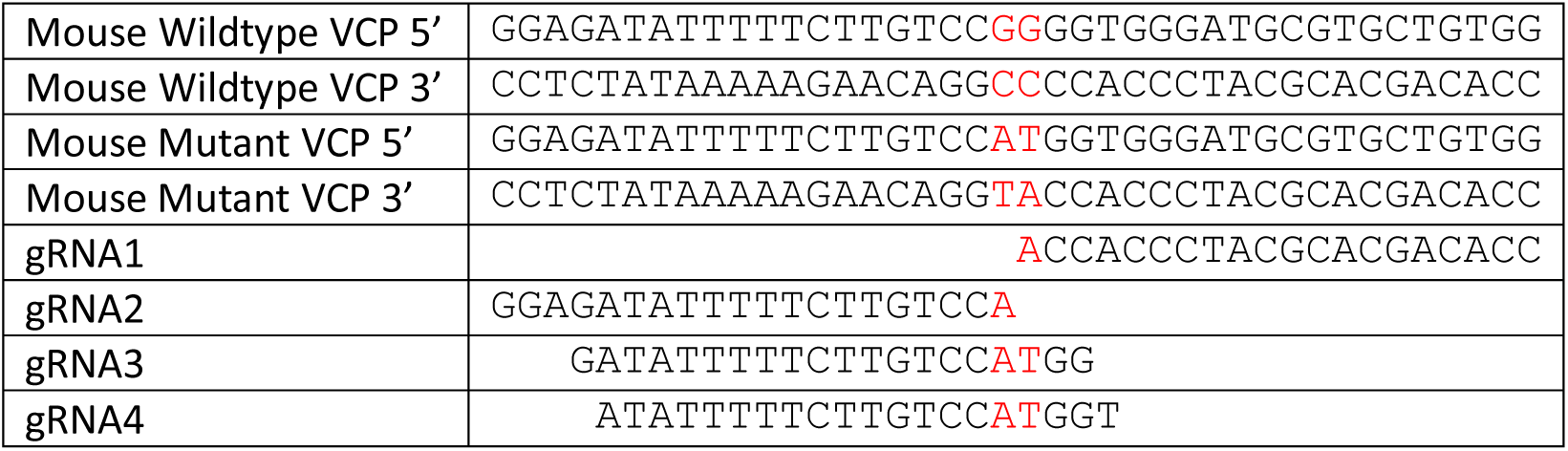
Targeting VCP c.464_465GG>AT in mouse myoblasts.

**Supplementary Figure 1.**
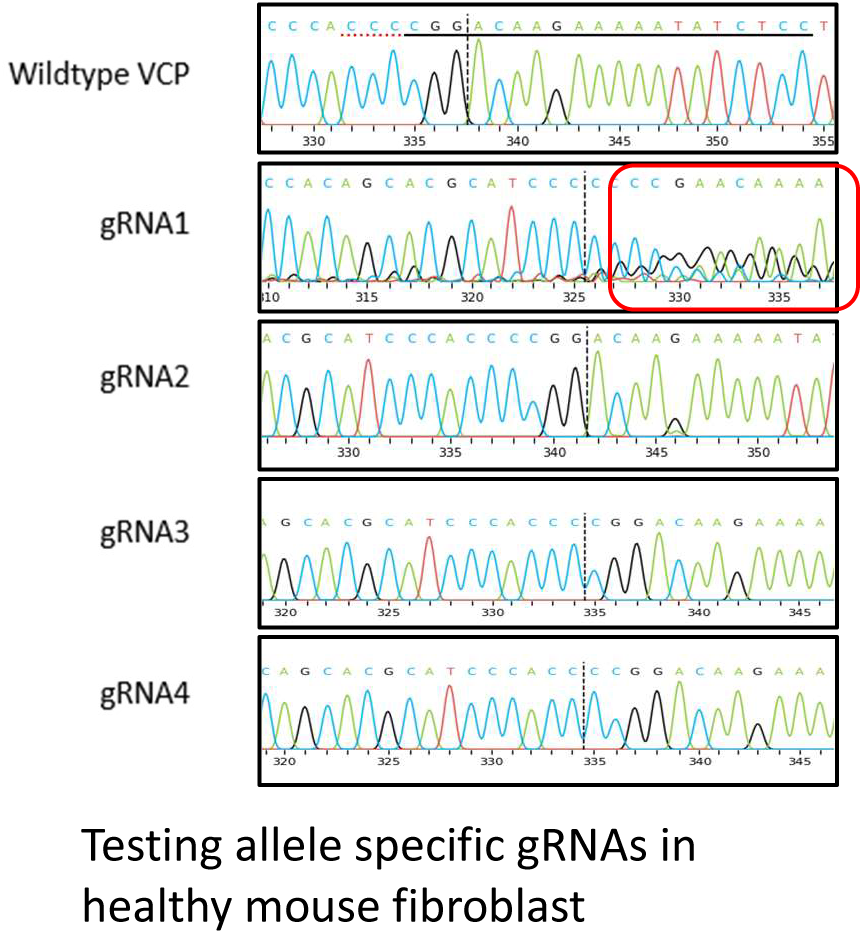
Testing REMEDY in mouse cells to detect on-target and off-targets. In PAM or near-PAM gRNAs (Supplementary table 1) strategies were tested in mouse healthy fibroblasts (Supplementary figure 1) and *VCP^R155H/+^* mouse myoblasts (Supplementary figure 2) to study on-target and off-targets measured by Sanger and NGS sequencing (Supplementary figure 3).

**Supplementary Figure 2.**
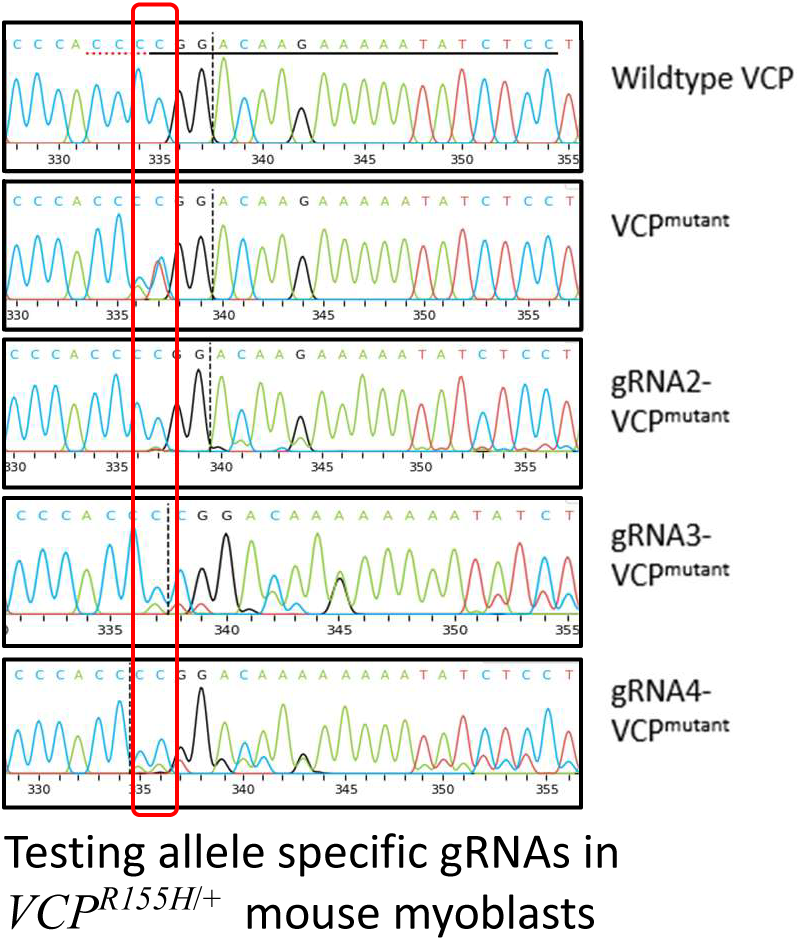
Testing REMEDY in mouse cells to detect on-target and off-targets. In PAM or near-PAM gRNAs (Supplementary table 1) strategies were tested in mouse healthy fibroblasts (Supplementary figure 1) and *VCP^RR155H/+^* mouse myoblasts (Supplementary figure 2) to study on-target and off-targets measured by Sanger and NGS sequencing (Supplementary figure 3).

**Supplementary Figure 3.**
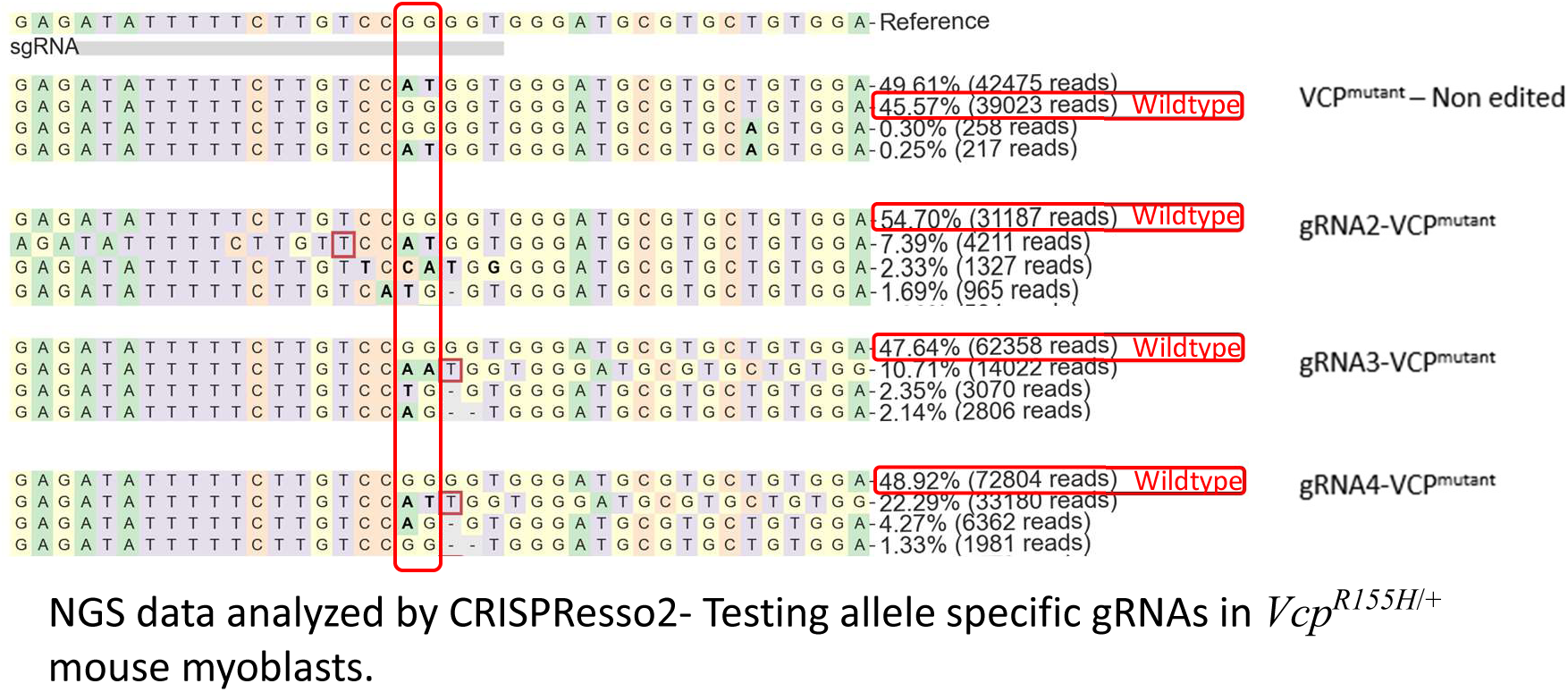
Testing REMEDY in mouse cells to detect on-target and off-targets. In PAM or near-PAM gRNAs (Supplementary table 1) strategies were tested in mouse healthy fibroblasts (Supplementary figure 1) and *VCP^R155H/+^* mouse myoblasts (Supplementary figure 2) to study on-target and off-targets measured by Sanger and NGS sequencing (Supplementary figure 3).

**Supplementary Figure 4.**
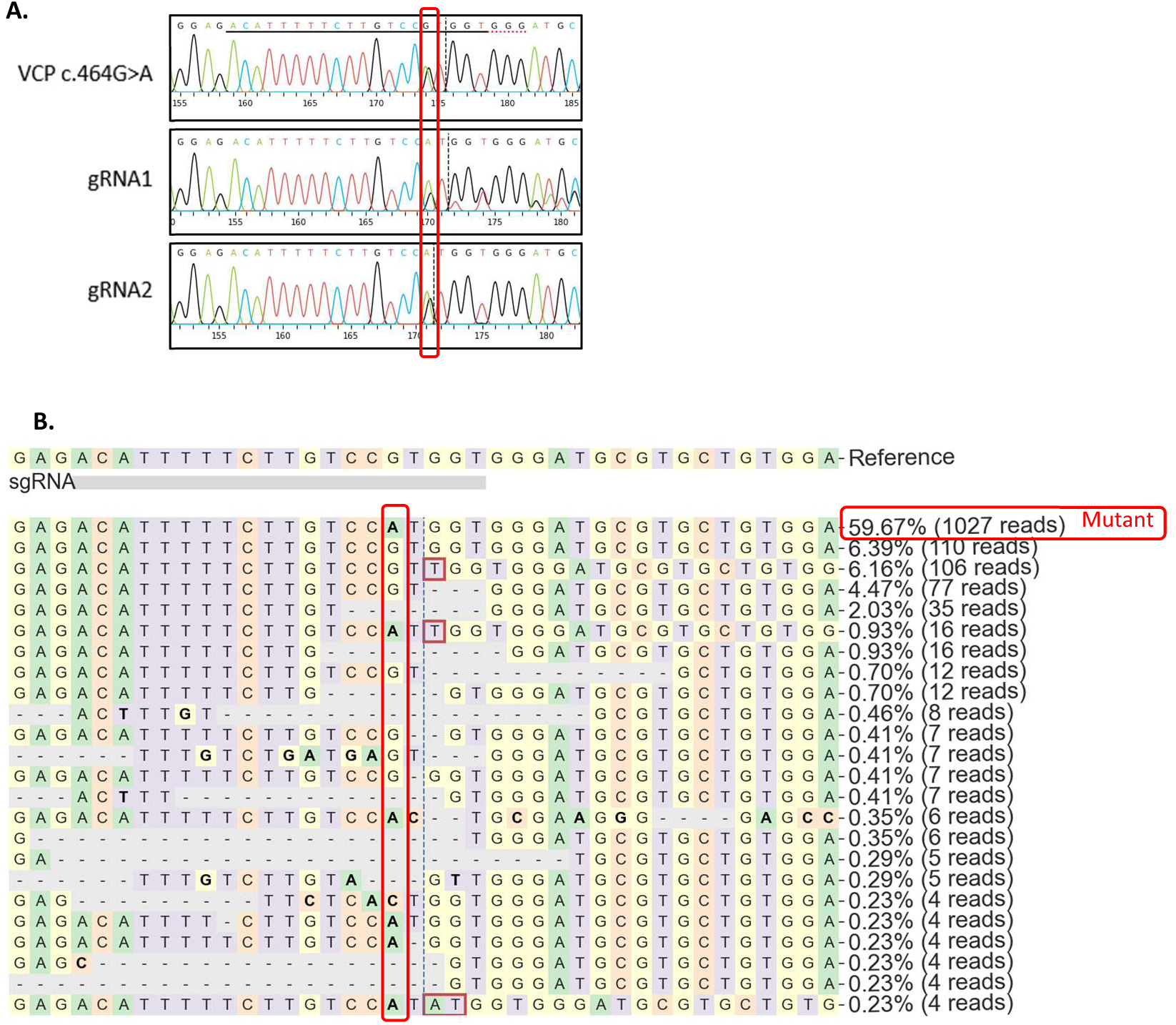
Testing REMEDY in c.464G>A (p.Arg155His) patient derived human fibroblast targeting wildtype allele, VCP measured by Sanger sequencing (A) and NGS (B).

**Supplementary Figure 5.**
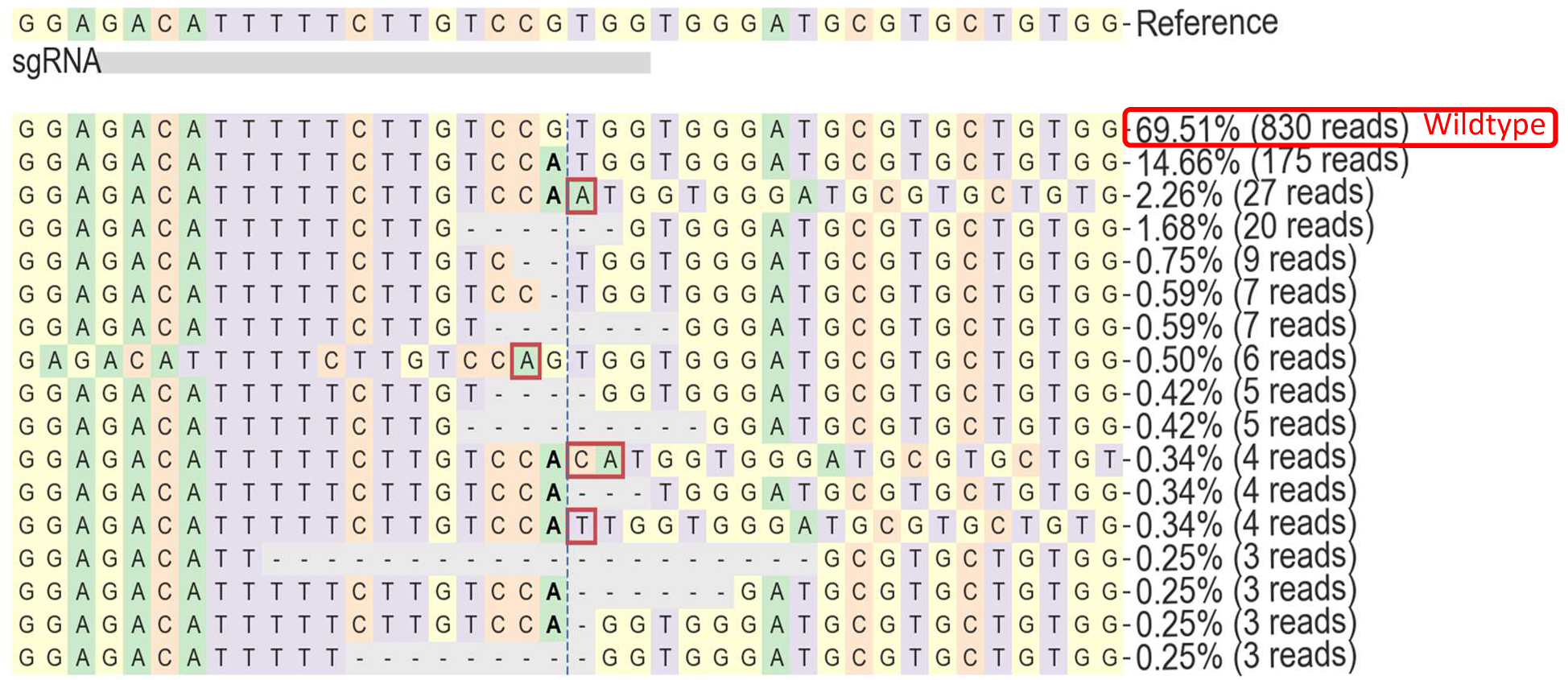
Testing REMEDY in c.464G>A (p.Arg155His) patient iPSC derived myoblasts targeting mutant allele, VCP measured by NGS.

**Supplementary Table 2.**
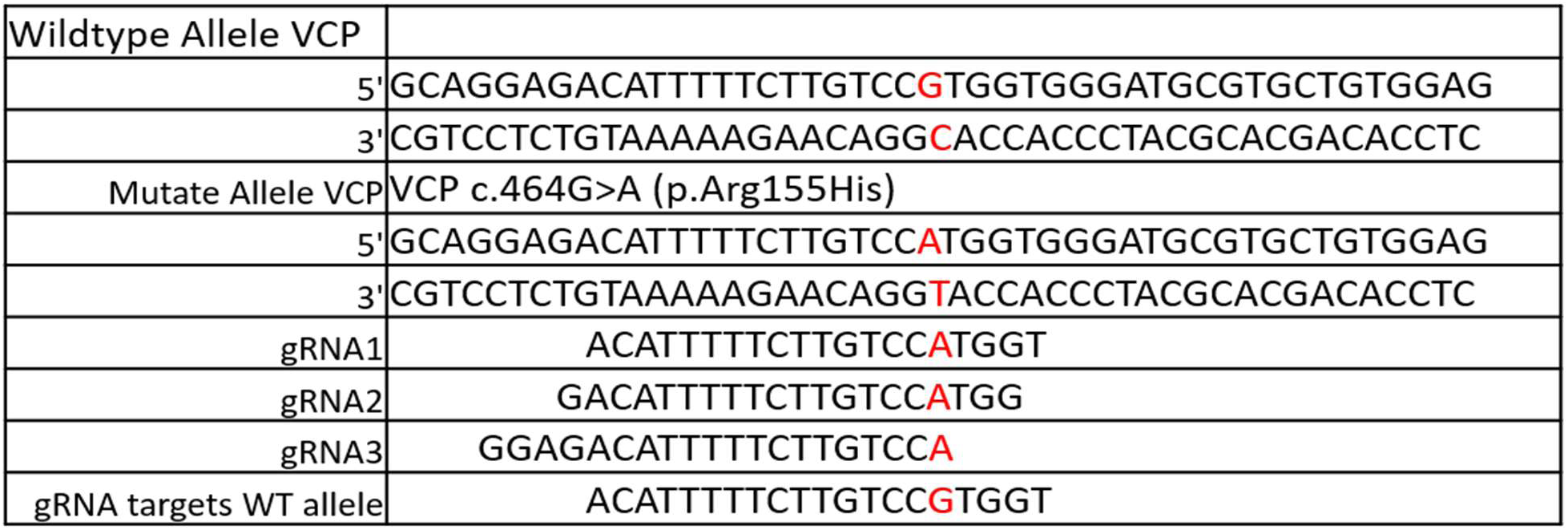
In-PAM or near-PAM gRNAs tested to target healthy and mutant allele in VCP c.464G>A (p.Arg155His) patient derived human fibroblasts.

**Supplementary Table 3.**
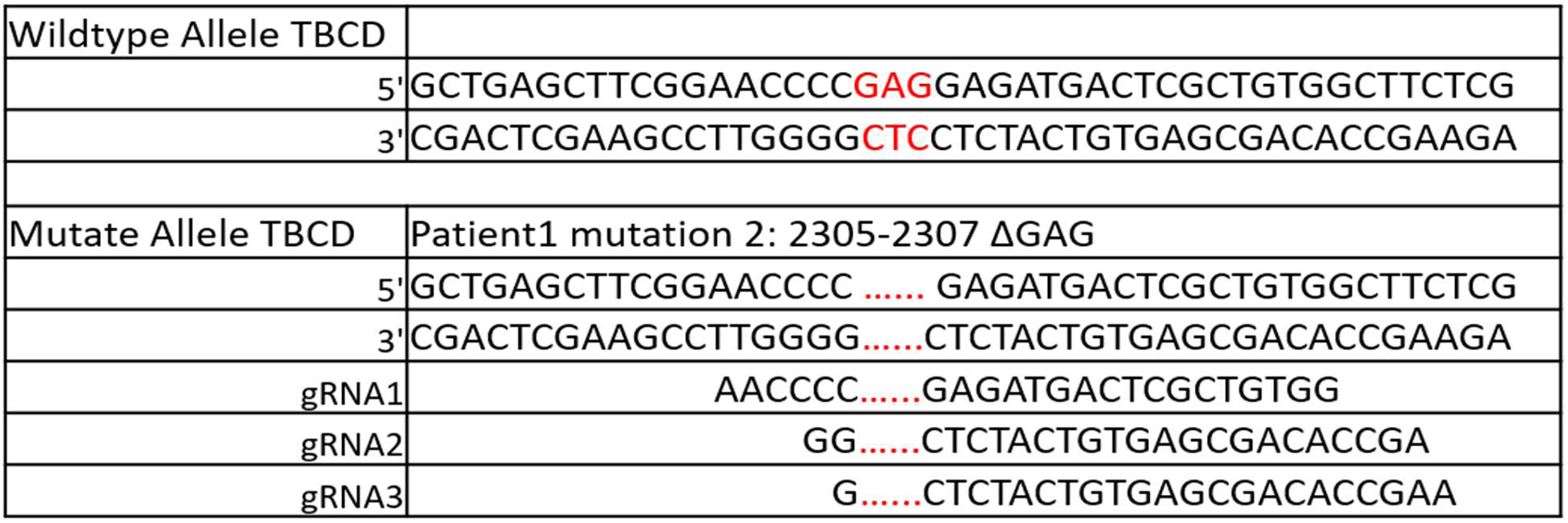
In-PAM or near-PAM gRNAs tested to target heterozygous mutation TBCD c.2305-2307 del GAG p.E769del (P1M2)

**Supplementary Table 4.**
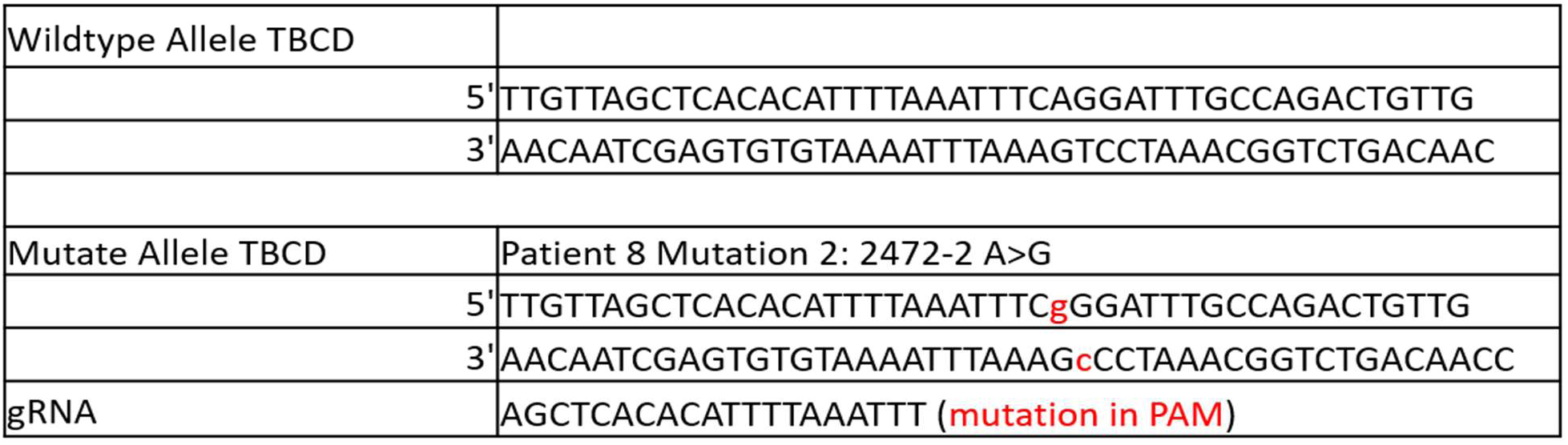
In-PAM or near-PAM gRNAs tested to target heterozygous mutation TBCD (P8M2)

**Supplementary figure 6.**
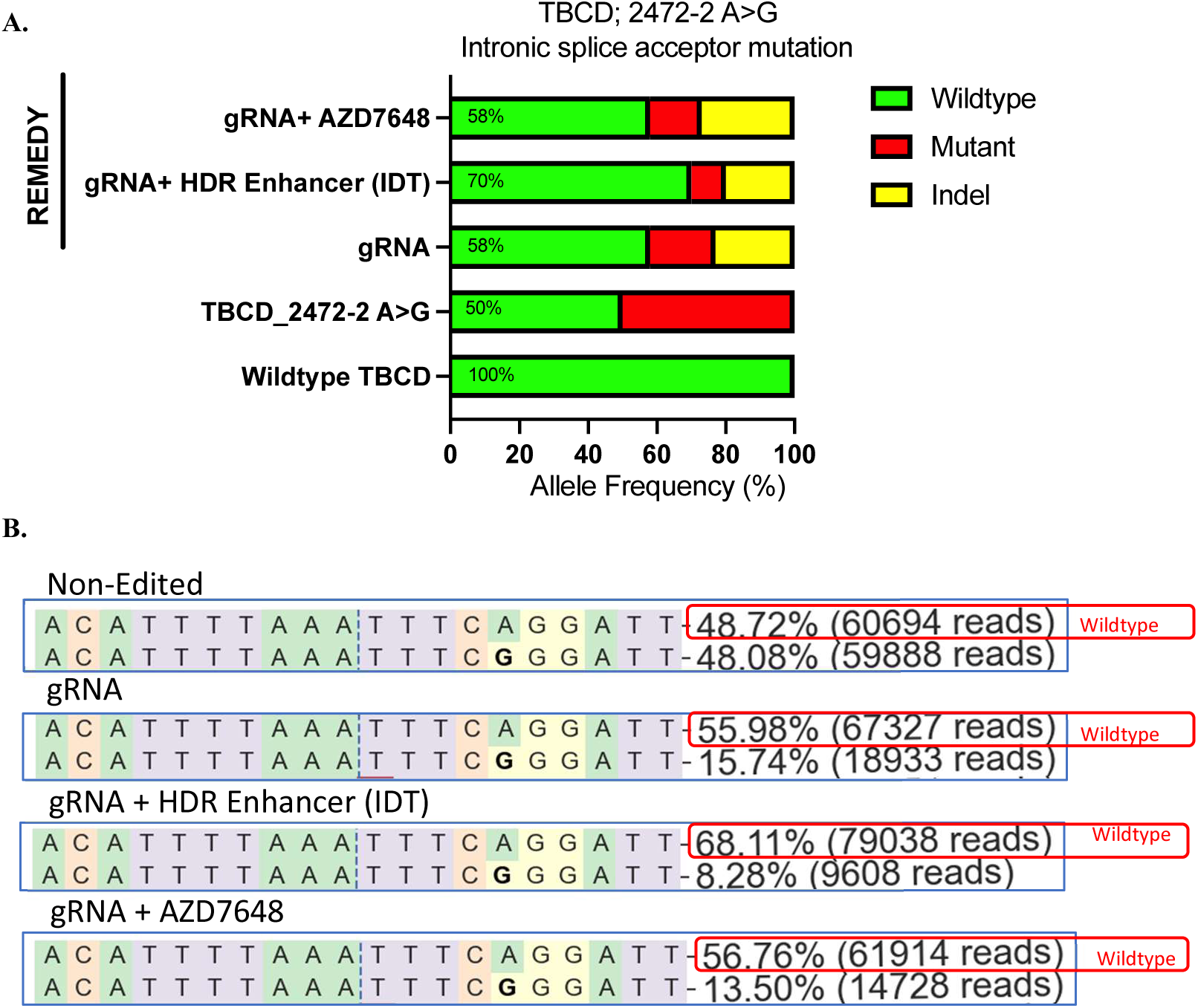
Testing REMEDY in human patient derived skin fibroblast with heterozygous mutation in TBCD (P8M2). Efficiency of REMEDY studied by deep Sanger sequencing (A) and NGS (B)

**Supplementary Table 5.**
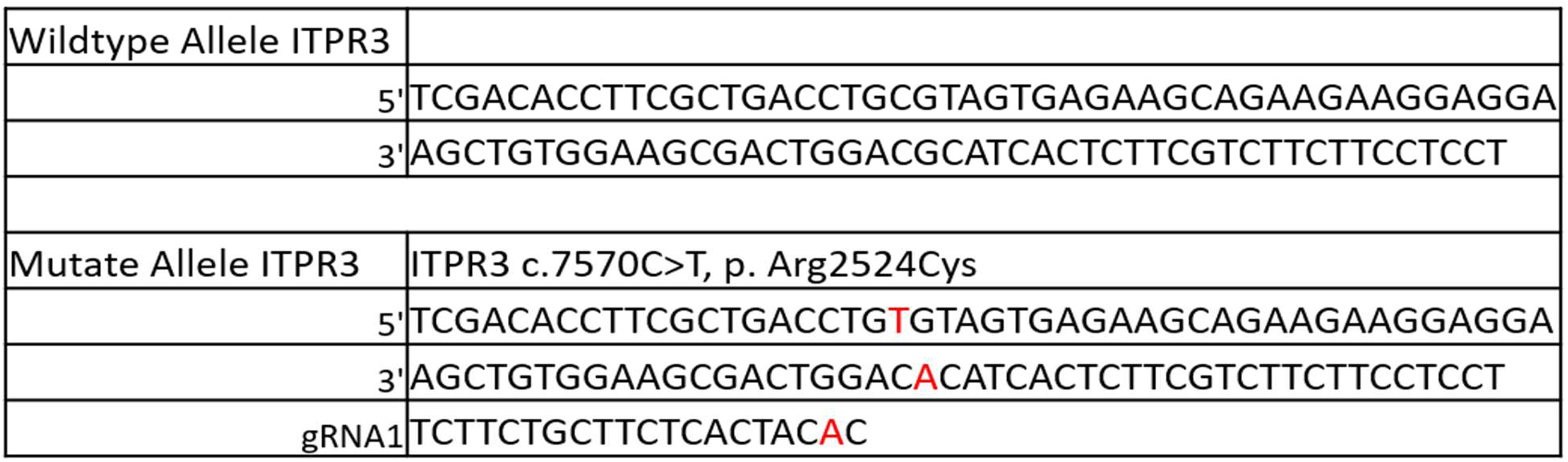
In-PAM or near-PAM gRNAs tested to target heterozygous mutation ITPR3 c.7570C>T, p. Arg2524Cys.

**Supplementary figure 8.**
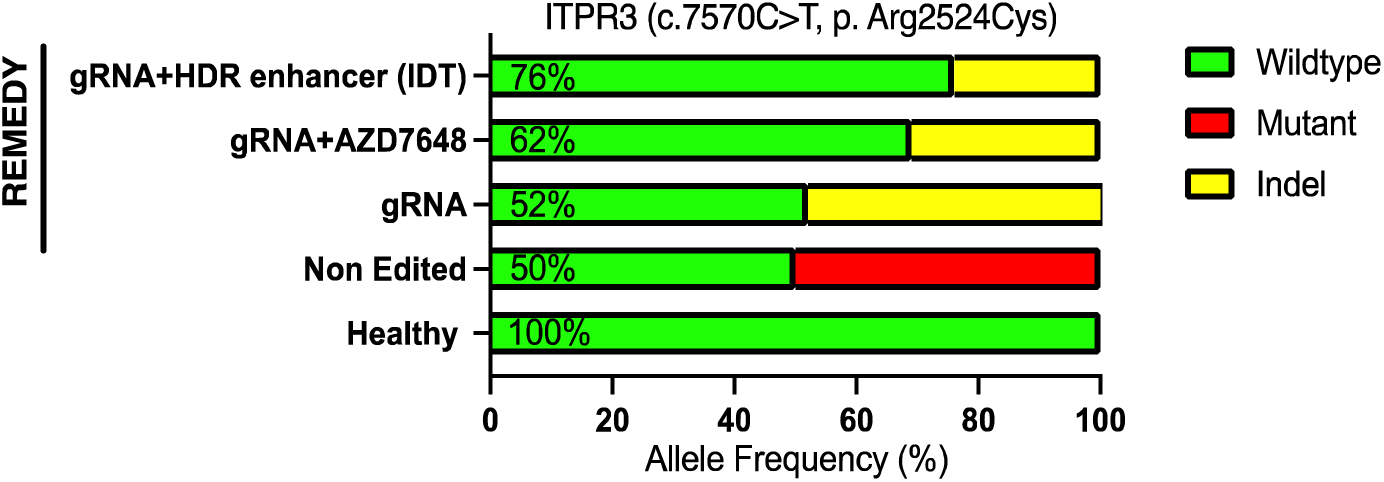
Sanger sequencing data of the *ITPR3* gene targeted in human skin fibroblast with heterozygous mutation. Data was analyzed by ICE.

**Supplementary Table 6.**
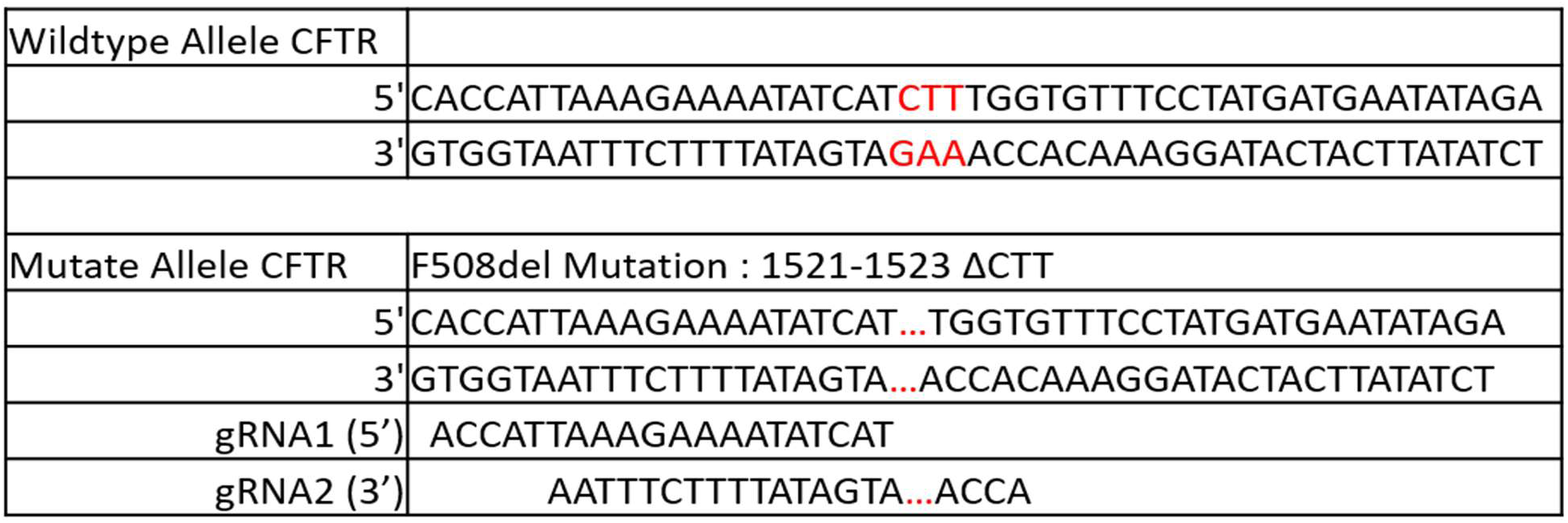
In-PAM or near-PAM gRNAs tested to target heterozygous mutation CFTR.

**Supplementary Table 7.**
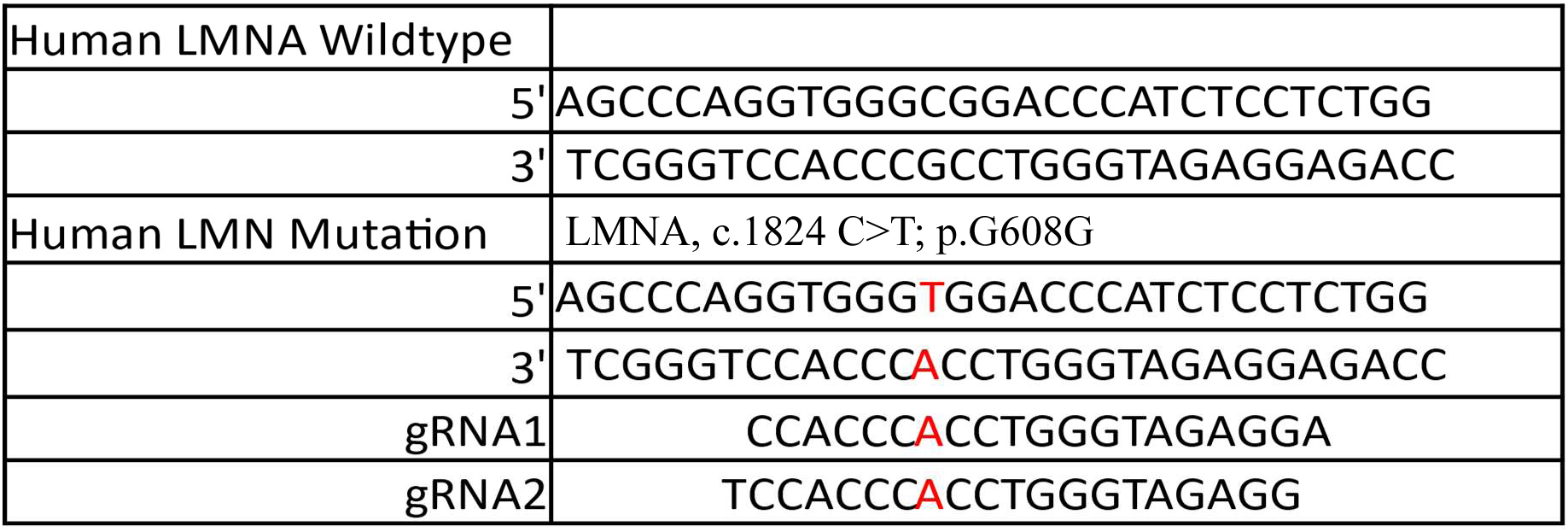
In-PAM or near-PAM gRNAs tested to target heterozygous mutation LMNA.

**Supplementary Table 8.**
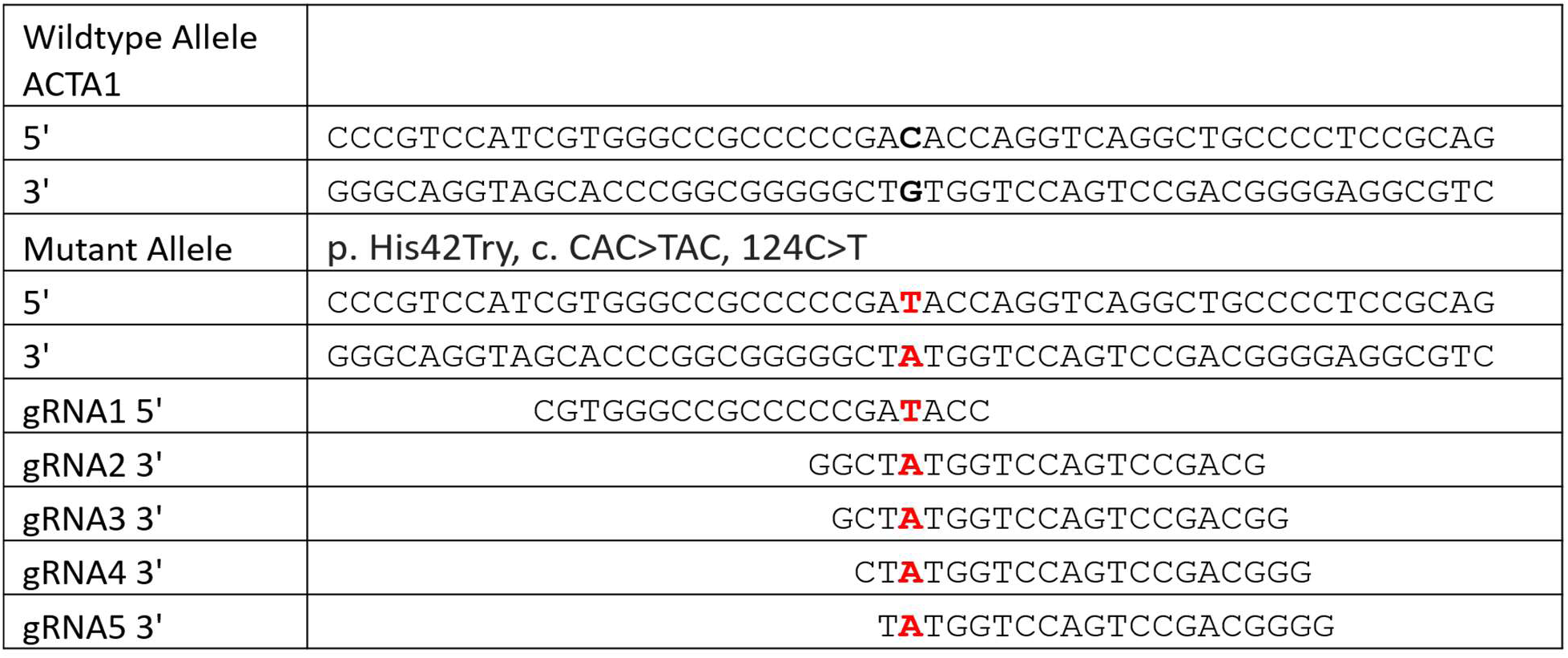
In-PAM or near-PAM gRNAs tested to target heterozygous mutation ACTA1.

**Supplementary Table 9.** Breakdown of alleles in Untreated and Post-REMEDY corrected samples. Counts and percentages are shown for each allelic category observed (A). Definitions of each of the 5 categories considered (B) are also shown. Categories can represent multiple allelic states.

| Type | Untreated |  | Post-REMEDY corrected |  |
| --- | --- | --- | --- | --- |
|  | Count (#) | Percent (%) | Count (#) | Percent (%) |
| Wild-Type | 20 | 52.6% | 22 | 56.4% |
| delGAG | 18 | 47.4% | 2 | 5.1% |
| Small In-Frame | 0 | 0 | 5 | 12.8% |
| Small Out-of-Frame | 0 | 0 | 8 | 20.5% |
| Large | 0 | 0 | 2 | 5.1% |
| Total | 38 | - | 39 | - |

Supplementary Table 9.
| Type | Description |
| --- | --- |
| Wild-Type | No deletion at chr17:82924983. Produces healthy, functional protein. |
| delGAG | Three-base in-frame deletion of Glutamic Acid, producing disease-causing protein. |
| Small In-Frame | 6-12 bp deletion without frameshift. |
| Small Out-of-Frame | <100bp deletion, resulting in a non-functional protein. |
| Large | >=100bp deletion, resulting in a non-functional protein. |

**Supplementary Table 10.** Phasing of Heterozygous Alleles Near Target Site. Reference and alternate allele counts are given for each of 33 phased heterozygous loci within 10kb of the target site. “delGAG In-Phase” indicates which allele is expected on reads containing a GAG deletion or any observed post-REMEDY deletion variant. Only 0.2% of reads contained an allele that deviated from expectations.

| Chrom. | Position | Ref. | Alt. | Untreated Ref. | Untreated Alt. | Corrected Ref. | Corrected Alt. | delGAG In-Phase | # Reads | # Deviated | % Deviation |
| --- | --- | --- | --- | --- | --- | --- | --- | --- | --- | --- | --- |
| chr17 | 82915798 | G | C | 16 | 24 | 12 | 19 | Ref | 31 | 0 | 0.0% |
| chr17 | 82916226 | A | C | 16 | 23 | 14 | 20 | Ref | 34 | 0 | 0.0% |
| chr17 | 82916414 | T | G | 17 | 22 | 14 | 20 | Ref | 34 | 0 | 0.0% |
| chr17 | 82916667 | G | T | 17 | 21 | 13 | 19 | Ref | 32 | 0 | 0.0% |
| chr17 | 82917553 | G | T | 24 | 16 | 20 | 15 | Alt | 35 | 0 | 0.0% |
| chr17 | 82917651 | A | G | 24 | 16 | 19 | 14 | Alt | 33 | 0 | 0.0% |
| chr17 | 82917750 | C | G | 16 | 21 | 14 | 20 | Ref | 34 | 0 | 0.0% |
| chr17 | 82918404 | C | T | 17 | 21 | 16 | 20 | Ref | 36 | 0 | 0.0% |
| chr17 | 82918537 | G | A | 16 | 21 | 15 | 20 | Ref | 35 | 0 | 0.0% |
| chr17 | 82918990 | T | A | 16 | 20 | 14 | 21 | Ref | 35 | 0 | 0.0% |
| chr17 | 82919236 | A | G | 15 | 20 | 14 | 20 | Ref | 34 | 0 | 0.0% |
| chr17 | 82919244 | G | T | 15 | 21 | 14 | 20 | Ref | 34 | 0 | 0.0% |
| chr17 | 82919556 | T | A | 15 | 21 | 14 | 20 | Ref | 34 | 0 | 0.0% |
| chr17 | 82919677 | C | T | 15 | 21 | 13 | 20 | Ref | 33 | 0 | 0.0% |
| chr17 | 82919986 | G | A | 14 | 21 | 16 | 20 | Ref | 36 | 0 | 0.0% |
| chr17 | 82920940 | C | T | 20 | 14 | 18 | 17 | Alt | 35 | 0 | 0.0% |
| chr17 | 82921116 | T | C | 14 | 20 | 17 | 18 | Ref | 35 | 0 | 0.0% |
| chr17 | 82924342 | T | C | 14 | 19 | 18 | 23 | Ref | 41 | 0 | 0.0% |
| chr17 | 82924487 | T | A | 16 | 19 | 17 | 21 | Ref | 38 | 0 | 0.0% |
| chr17 | 82927573 | G | C | 25 | 15 | 20 | 16 | Alt | 36 | 0 | 0.0% |
| chr17 | 82928440 | G | T | 17 | 23 | 18 | 20 | Ref | 38 | 0 | 0.0% |
| chr17 | 82929368 | T | C | 18 | 22 | 20 | 19 | Ref | 39 | 0 | 0.0% |
| chr17 | 82929575 | A | G | 18 | 21 | 20 | 19 | Ref | 39 | 0 | 0.0% |
| chr17 | 82930288 | A | G | 16 | 22 | 20 | 19 | Ref | 39 | 0 | 0.0% |
| chr17 | 82931029 | T | C | 15 | 22 | 21 | 22 | Ref | 43 | 0 | 0.0% |
| chr17 | 82932386 | C | T | 17 | 22 | 20 | 20 | Ref | 40 | 0 | 0.0% |
| chr17 | 82932762 | T | C | 19 | 21 | 20 | 18 | Ref | 38 | 2 | 5.3% |
| chr17 | 82933259 | C | T | 20 | 21 | 21 | 19 | Ref | 40 | 0 | 0.0% |
| chr17 | 82933517 | A | G | 19 | 21 | 19 | 18 | Ref | 37 | 0 | 0.0% |
| chr17 | 82933584 | C | T | 20 | 20 | 19 | 18 | Ref | 37 | 0 | 0.0% |
| chr17 | 82933790 | G | T | 20 | 20 | 19 | 18 | Ref | 37 | 0 | 0.0% |
| chr17 | 82933799 | T | G | 20 | 20 | 19 | 18 | Ref | 37 | 0 | 0.0% |
| chr17 | 82933802 | A | G | 20 | 20 | 18 | 17 | Ref | 35 | 0 | 0.0% |
| Total | - | - | - | 581 | 671 | 566 | 628 | - | 1194 | 2 | 0.2% |

**Supplementary Figure 9.**
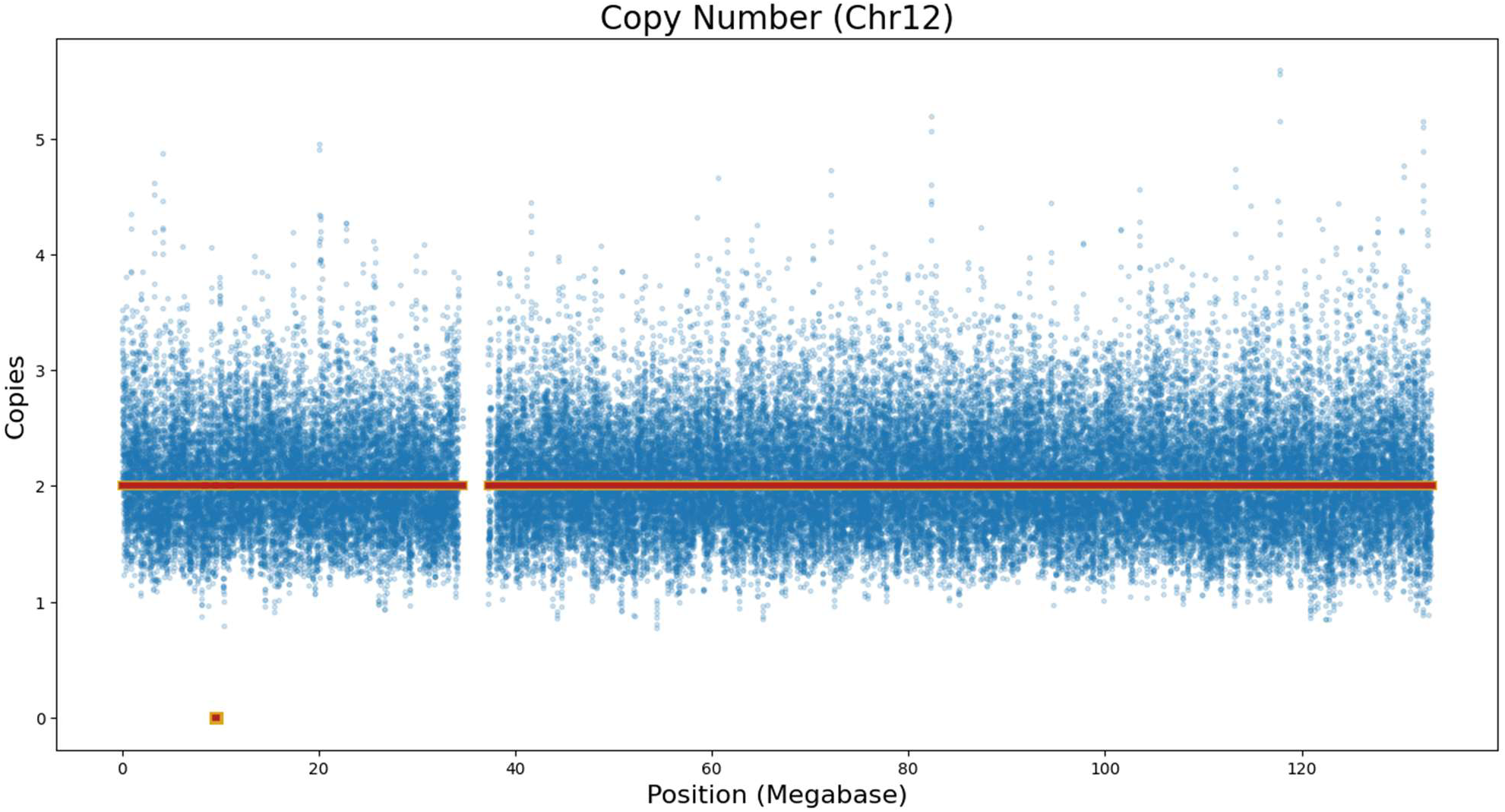
Copy-Number Analysis of Chromosome 17. Blue background points show copy number calculated at a given location from post-treatment coverage depth, normalized against coverage of the pre-treatment sample. Yellow and red segments show the HiFiCNV-calculated copy state at a given position for the preand post-treatment samples, respectively. Overlapping red and yellow segments show copy states which

**Supplementary Figure 10.**
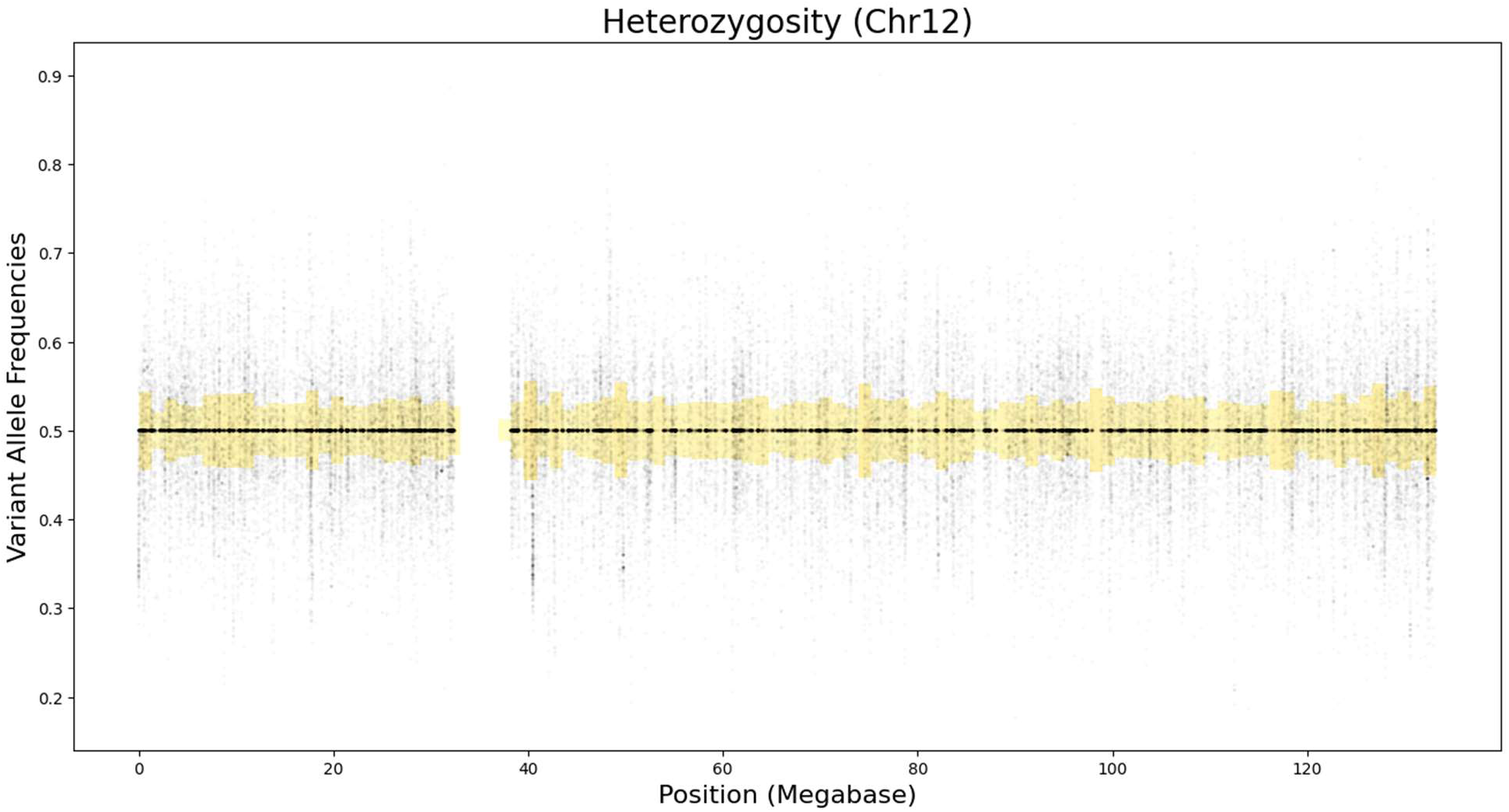
Loss-of-Heterozygosity Analysis of Chromosome 12. Black points show variant allele frequency called at a given position. Yellow bars show allelic spread (symmetric distance of calculated average from 0.5) for regions each representing 1% of the chromosome.

**Supplementary Table 11.**
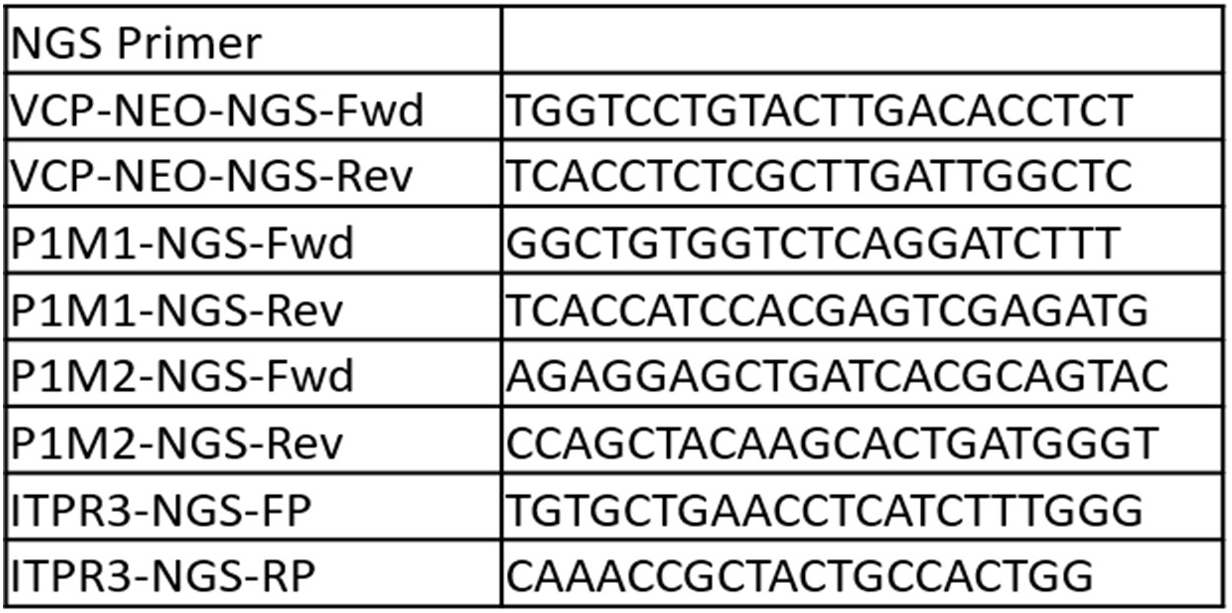
PCR Primers used for Next Generation Sequencing.

**Supplementary Table 12.**
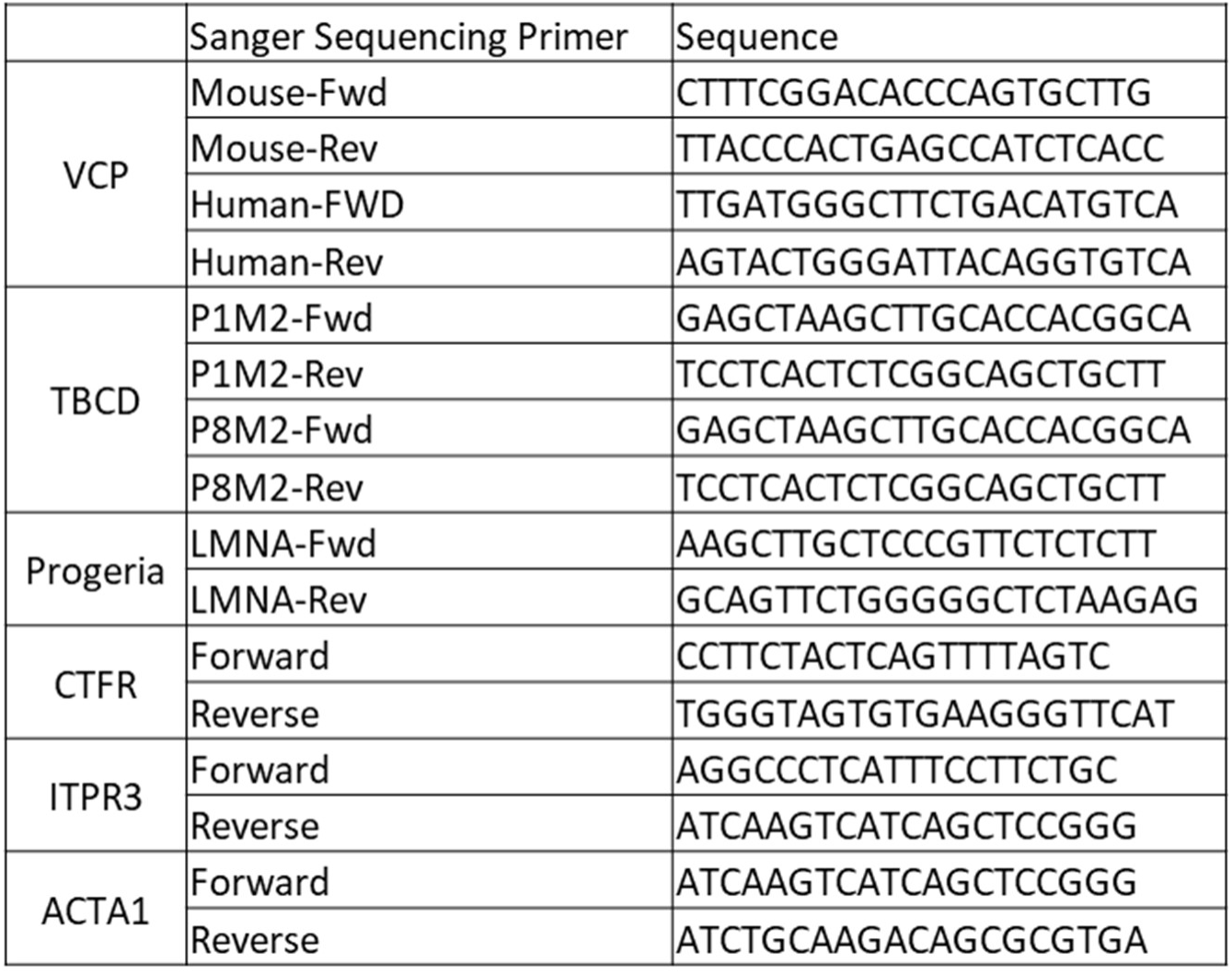
PCR Primers used for Sanger sequencing.

